# Genetic Ablation and Multi-Omics Profiling Reveal CEP55 as a Key Driver of Tumorigenesis in Diverse Cancer Models

**DOI:** 10.1101/2025.10.28.685219

**Authors:** Behnam Rashidieh, Yuanhao Yang, Amanda Louise Bain, Phillip Theron, Catharina Vosloo, Parinaz Ahangar, Brydie Bowden, Simon M. Tria, Ben B. Wang, Sandra Isenmann, Nicholas J. Westra van Holthe, Zherui Xiong, Hani Vu, Sowmya Sharma, Asmerom Sengal, J. Alejandro Lopez, John Finnie, Antonella Papa, Pascal HG Duijf, Quan Nguyen, Pirjo M Apaja, Kum Kum Khanna

## Abstract

**Background:** CEP55 is a centrosomal protein with emerging oncogenic roles. It is overexpressed in many cancers, where it drives genomic instability, phenocopies PTEN loss and hyperactivates the PI3K/AKT pathway, thereby promoting aggressive and therapy-resistant tumors.

**Methods:** We developed an inducible Cep55 knockout (KO) mouse model as well as generated primary, immortalized (SV40), and transformed (E1A/Ras) mouse embryonic fibroblasts (MEFs) for functional and proteomic assays. Tumorigenesis was studied using E1A/Ras MEF transplantation and a *Pten-*deficient genetically engineered mouse model (GEMM). Multi-omics apparatus including proteomics, and spatial transcriptomics, histopathology, immunobloting and functional assays were employed to elucidate underlying mechanisms. Findings were cross-validated using human cancer datasets.

**Results:** Cep55 ablation remodeled the extracellular matrix (ECM), disrupted integrin and oncogenic signaling, reduced AKT/ERK activation, and induced stress pathways. Cep55 knockouts (KO) cells showed impaired proliferation, migration, invasion, and adhesion*. In vivo*, unlike the embryonic lethality observed in constitutive KOs, the inducible *Cep55* deletion was well-tolerated in adult mice. Strikingly, tumors derived from *Cep55*-null EIA/RAS-transformed MEFs exhibited delayed onset and progression. In the genetically engineered cancer-prone mouse model, *Cep55* deletion on a *Pten*-deficient background led to a greater than seven-fold increase in median survival. Spatial transcriptomics on cancerous tissues revealed that CEP55 regulates ECM remodeling, integrin expression, trafficking, and cell adhesion, supporting its critical role in tumor progression to metastasis.

**Conclusion:** CEP55 is dispensable for adult homeostasis but essential for tumorigenesis, especially with PTEN loss. Its deletion suppresses transformation, delays tumor growth, and prolongs survival, supporting CEP55 as a therapeutic target in PTEN-deficient cancers.

**Graphical Abstract:** 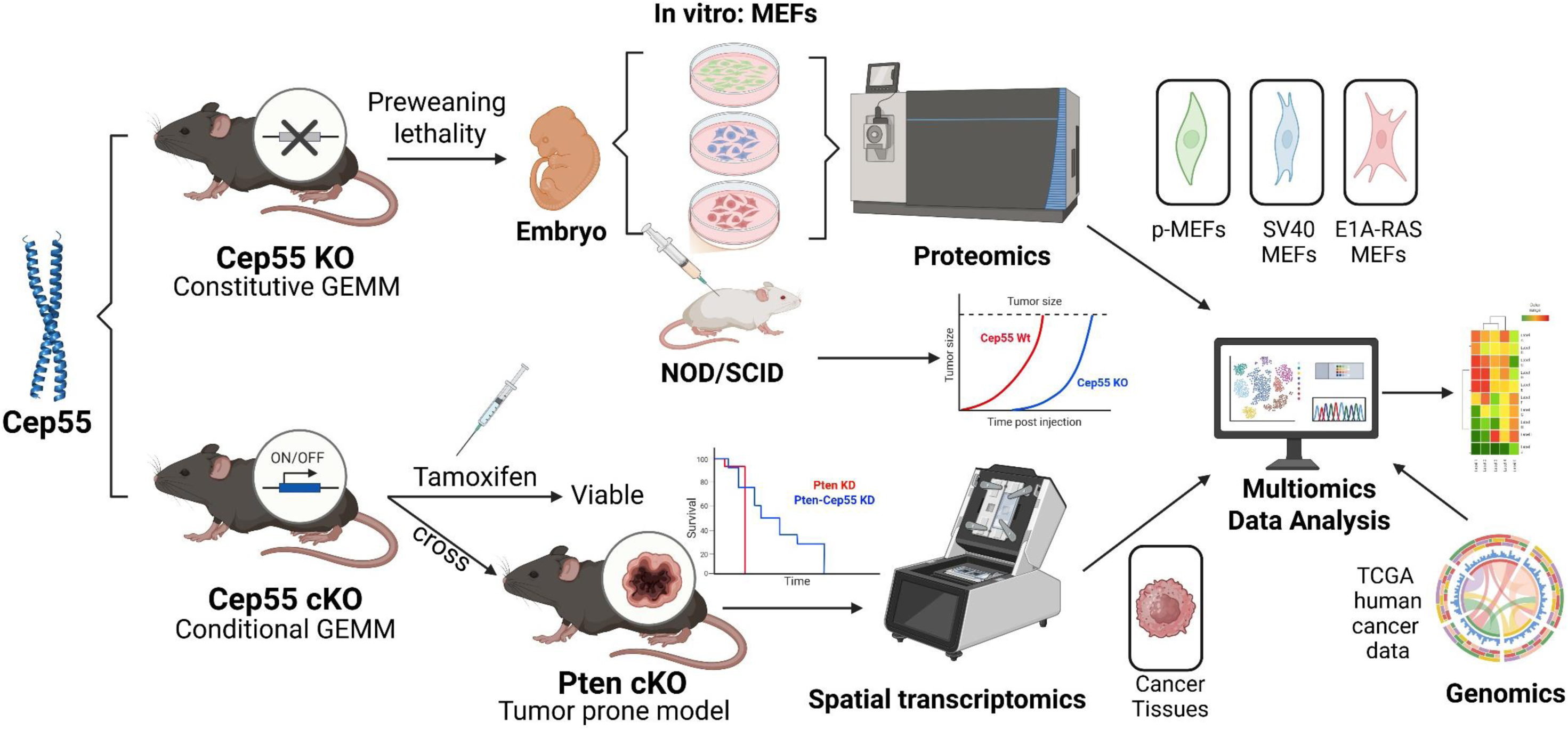

## Background

Centrosomal protein 55 (CEP55) plays a crucial role in cell division, particularly during the final stage of cytokinesis. It is recruited to the midbody of dividing cells, where it facilitates the recruitment of other essential proteins, such as endosomal sorting complexes required for transport (ESCRT) machinery, to ensure the proper separation of daughter cells (Fabbro et al., 2005, Zhao et al., 2006). Given its fundamental role in regulation of cell proliferation, it is not surprising that CEP55 dysregulation has been implicated in the development and progression of numerous human cancers. CEP55 is overexpressed across a wide range of malignancies, including breast, lung, colon, liver cancers, T-cell lymphoma and is frequently associated with poor prognosis, advanced disease stage, metastasis and resistance to immunotherapy (Jeffery et al., 2016). While normally silenced in adult tissues, CEP55 is aberrantly reexpressed in many cancers and is classified as a cancer-testis antigen. Functionally, it promotes oncogenic signaling by interacting with PIK3CA leading to enhanced PI3K/AKT pathway activity (Chen et al., 2007).Through this mechanism, CEP55 can hyperactivate AKT-driven survival pathways and promote unchecked cell proliferation (Rashidieh et al., 2021b, Sinha et al., 2018, Sinha et al., 2020).

Overexpression of CEP55 alone is sufficient to induce tumorigenesis in a genetically engineered mouse model and is associated with activation of PI3K/AKT pathway, mitotic defects and genomic instability (Sinha et al., 2020). Conversely, CEP55 knockdown or silencing impairs cancer cell proliferation and viability, inducing apoptosis. These findings position CEP55 as an oncogenic driver and an attractive candidate for targeted cancer therapy.

In contrast, PTEN (the phosphatase and tensin homolog) is one of the most frequently inactivated tumor suppressors in human cancer (Lee et al., 2018). PTEN antagonizes the PI3K/AKT pathway by dephosphorylating PIP3 to PIP2, thereby inhibiting AKT activation and downstream proliferative signaling. Loss of PTEN function via mutations, genomic deletion, or epigenetic silencing results in constitutive activation of PI3K/AKT signaling, promoting enhanced cancer cell survival, proliferation, angiogenesis and invasiveness. Biallelic PTEN Loss (deletions or mutations) are observed in numerous malignancies, including aggressive subsets of breast cancer, prostate, glioblastoma, and endometrial carcinoma (Hollander et al., 2011). In mouse models, systemic heterozygous *Pten* deletion drives the rapid development of multifocal tumors (notably lymphomas and carcinomas), underscoring its broad tumor-suppressive function (Borlado et al., 2000, Stahl et al., 2003). Clinically, PTEN-deficient tumors are associated with poor outcomes and frequent resistance to PI3K/AKT/mTOR inhibitors (Browne and Okines, 2024, Mao et al., 2025). PTEN loss fosters an immunosuppressive tumor microenvironment, reducing responses to immune checkpoint blockade therapies (Peng et al., 2016). These challenges underscore the urgent need for alternative therapeutic strategies in PTEN-deficient cancers (Vidotto et al., 2020).

Notably, oncogenic activity of CEP55 and the effects of PTEN loss converge on the same pro-survival PI3K/AKT axis. CEP55-driven PI3K stabilization and AKT activation functionally mimic the hyperactivation of this pathway caused by PTEN loss. In PTEN-deficient cancers, CEP55 may become particularly essential for sustaining elevated PI3K/AKT signaling. We therefore hypothesized that disrupting CEP55 in a PTEN-deficient context could attenuate tumor progression by restoring suppression of AKT signaling, potentially succeeding where direct pathway inhibitors have failed.

In this study, we developed a novel tamoxifen-inducible Cep55 conditional knockout mouse and evaluated the impact of Cep55 loss on cellular transformation and tumorigenesis both *in vitro* and *in vivo*. This included studies in genetically engineered mouse models combining inducible *Cep55 deletion* with *Pten* loss. Specifically, we investigated whether dual Cep55-Pten knockdown (CP DKD) would hamper tumorigenesis in mice with loss of Pten alone. Our findings provide crucial insights into CEP55’s role in cancer development and suggest that targeting CEP55 may represent a viable strategy in PTEN-deficient cancers.

## Methodology

### Mouse Models

A *Cep55* floxed allele was engineered (knockout-first allele construct) and crossed with Rosa26-Cre^ERT2^ mice to generate an inducible knockout (Cep55^Fl/Fl^; Cre^ERT2+/−^). Tamoxifen (single intraperitoneal dose, 1 mg) was administered to 6–8-week-old mice to induce global Cep55 deletion (cKO). A *Pten*^Fl/Fl^ line was crossed to Cep55 cKO mice to produce Cep55/Pten double conditional knockouts. Mice were monitored for health for up to 160 days post-induction of tamoxifen. Stable Cep55 knockout was confirmed by PCR and immunoblot. All animal procedures were approved by institutional ethics committees, QIMR Berghofer Animal Ethics Committee (A0707-606M).

### Tumor-Prone Pten loss Model

Cep55^Fl/Fl^; Pten^Fl/Fl^; Cre^ERT2+/−^ mice and Pten^Fl/Fl^; Cre^ERT2+/−^ only controls (6–8 weeks old) were induced with tamoxifen to delete both genes (Cep55-Pten DKD) or Pten KD alone. Mice were monitored for morbidity. Moribund or ≥160 days post-injection survivors (3) were necropsied. Tissues (thymus, lymph nodes, spleen, liver, etc.) were examined by a veterinary pathologist. Histopathological analysis, including Hematoxylin and Eosin (H&E) staining and immunohistochemistry (IHC), was performed as previously described (Rashidieh et al., 2023) on tumor tissues to assess tumor morphology and protein expression. Tumors were classified and incidence recorded. Kaplan–Meier survival curves were generated for each cohort.

### Transplant cancer cell-derived Model

10^6^ E1A/Ras-transformed MEFs (Cep55 Wt or KO) 1:1 in Matrigel were injected subcutaneously into NOD/SCID (Non-Obese Diabetic/Severe Combined Immunodeficiency) mice (n=6 per group). Mice were examined thrice weekly for palpable tumors and sacrificed at endpoint tumor size (∼1000 mm^3^). Tumor volume was measured by calipers. Kaplan–Meier analysis compared tumor-free survival between groups. Harvested tumors were formalin-fixed for histology.

### Cell Culture and Transformation

Primary mouse embryonic fibroblasts (pMEFs) were isolated from E13.5 Cep55^+/+^ and Cep55^−/−^ embryos (the latter obtained from heterozygous intercrosses). MEFs were cultured in DMEM + 20% FBS and either used as primary cells or immortalized by SV40 large T antigen, or transformed by adenoviral E1A and oncogenic Ras (Ras^V12^) to model stepwise tumorigenesis using lentviral packaging system.

### Single Vesicular pH trafficking microscopy

Integrin and other cargo tracking using pH-sensitive fluorophores, as well as analysis, was performed as (Kazan et al., 2019) and (Kharitidi et al., 2015). ALIX minimal Cep55-interacting peptide (sequence GPPSLNY) (Lee et al., 2008) and control (GPASLNA incorporating alanine substitution) were synthesized with N-terminal myristylation and C-terminal amidation to enhance membrane association and peptide stability. Cells were treated with 20 µM peptides in serum free medium before the experiment. Live-cell image acquisition was carried out Nikon TI-E inverted microscope, using triggered acquisition, and analyzed with NIS-Elements (Nikon, Japan).

### Cellular Functional Assays

Proliferation was measured in real-time (IncuCyte live-cell imaging) over 5–7 days as previously described (Rashidieh et al., 2021a). Colony formation was assessed in 2D (plating 500 cells/well, 14 days) and 3D soft agar and Happy Cell suspension medium assays (14 days). Wound-healing migration assays and Transwell invasion assays (Matrigel-coated chambers) were performed to evaluate cellular plasticity. An adhesion assay measured cell attachment to collagen-coated plates. A custom invasion assay was established using a polyisocyanopeptide (PIC)-based hydrogel modified with the collagen mimetic peptide GFOGER (PIC-GFOGER) within ultra-low attachment plates (Westra van Holthe et al., 2024).

### Proteomic and Phosphoproteomic

Quantitative mass spectrometry was performed on Wt vs. Cep55-KO MEFs (primary, SV40-immortalized, and E1A/Ras-transformed) as previously described (Rashidieh et al., 2023). Total protein lysates were trypsin-digested and analyzed by high-resolution LC-MS/MS. Phosphopeptides were enriched by TiO_2_ chromatography. Over 2,000 proteins and ∼500 phosphosites were quantified per comparison.

### Spatial Transcriptomics

To investigate gene expression changes in splenic lymphomas, we performed 10x Genomics Visium spatial transcriptomics on FFPE spleen sections from Pten-KD and Cep55-Pten-DKD mice (n = 3 tumors per group). FFPE tissue sections were processed following the manufacturer’s instructions (Visium CytAssist Tissue Preparation Guide, CG000518). Briefly, 5 μm tissue sections mounted on glass slides were stained with hematoxylin and eosin (H&E) and imaged using the Aperio XT Brightfield Automated Slide Scanner (Leica, Germany) at 20× magnification. After imaging, tissue sections were destained and decrosslinked, then hybridized with mouse transcriptome probes using the 10x Genomics Mouse Transcriptome 6.5Lmm kit (PN #1000521, #1000365).

Probe extension and library construction were performed according to the standard Visium CytAssist FFPE workflow (Visium Spatial Gene Expression User Guide, CG000495). The libraries were sequenced using an Illumina NextSeq 2000 (Illumina, USA) with paired-end dual indexing: 28 cycles for Read 1, 10 cycles for i7, 10 cycles for i5, and 50 cycles for Read 2. Sequencing data were demultiplexed using bcl2fastq v2.20.0.422 (Illumina). Reads were aligned and mapped to the H&E images using Space Ranger v2.1.1 (10x Genomics) and the mm10 mouse reference genome (refdata-gex-mm10-2020-A). Following quality control to exclude low-quality spots using the Seurat v5 pipeline, differential gene expression analysis was performed to identify genes and pathways upregulated or downregulated upon Cep55 loss in the PTEN-KD tumor context. Analyses were conducted both across the whole TMA and within pathologist-annotated lymphoma regions.

### Immune Profiling, Genomics and Bioinformatics

Public TCGA RNA-seq data (32 cancer types) were mined for CEP55 expression correlations with ∼1,387 oncogenic signatures (cell cycle, mitosis, DNA repair, etc.) and immune cell infiltration scores (TIMER). Immune cell abundances were inferred by CIBERSORT and TISIDB, correlating CEP55 expression or copy-number with 28 tumor-infiltrating lymphocyte (TIL) types. TISIDB was further used to examine CEP55 associations with immunomodulators (checkpoint molecules, cytokines) and clinical outcomes across cancers. Differentially expressed mRNAs, proteins and phosphosites (Log2 foldchange >0.5, p<0.05) were subjected to Gene Ontology (GO) enrichment analysis to identify impacted pathways. All data analysis used R with appropriate packages; Pearson correlation, statistical significance was defined as p<0.05, in proteomics pathway clusters p<0.01.

### Statistical analysis

Two-tailed unpaired or paired Student’s t-test, one-way or two-way ANOVA with Bonferroni post hoc correction and log-rank testing were performed where appropriate using Prism v9.0 (Graph Pad Software, La Jolla, CA, USA) and the P-values were calculated as stated in the Figure legends. Statistical significance was noted to each Figure as follows: “no significance” (ns), P > 0.05; *P ≤ 0.05; **P ≤ 0.01, ***P ≤ 0.001, and ****P ≤ 0.0001. Mean ± SD: Mean and standard error of the mean were used to describe the variability within the sample in our analysis.

## Results

### CEP55 Is broadly overexpressed in human cancers and correlates with poor prognosis

Analysis of *CEP55* expression across The Cancer Genome Atlas (TCGA) dataset revealed that CEP55 is broadly overexpressed in a wide range of human malignancies, consistent with previous report (Zhou et al 2024). Pan-cancer RNA-seq analyses demonstrated significantly elevated CEP55 mRNA in the majority of tumor types when compared to their corresponding normal tissues. Notably, high expression levels were observed in breast invasive carcinoma (BRCA), bladder urothelial carcinoma (BLCA), colon adenocarcinoma (COAD), glioblastoma multiforme (GBM), lung adenocarcinoma (LUAD), lung squamous cell carcinoma (LUSC), ovarian serous cystadenocarcinoma (OV), and pancreatic adenocarcinoma (PAAD) (**Sup fig 1a**). Clinically, elevated CEP55 expression was associated with more aggressive disease characteristics, including advanced tumor stage and grade, increased incidence of metastasis and significantly shorter overall survival (hazard ratios> 1; log-rank p < 0.01) across multiple cancer types. Moreover, CEP55 expression increased progressively from stage I to IV and from low- to high-grade tumors.These associations were consistent across datasets and indicate that CEP55 upregulation is a strong marker of poor prognosis (**Sup fig 1b, Sup material_1_Fig a**).

**Figure 1:**
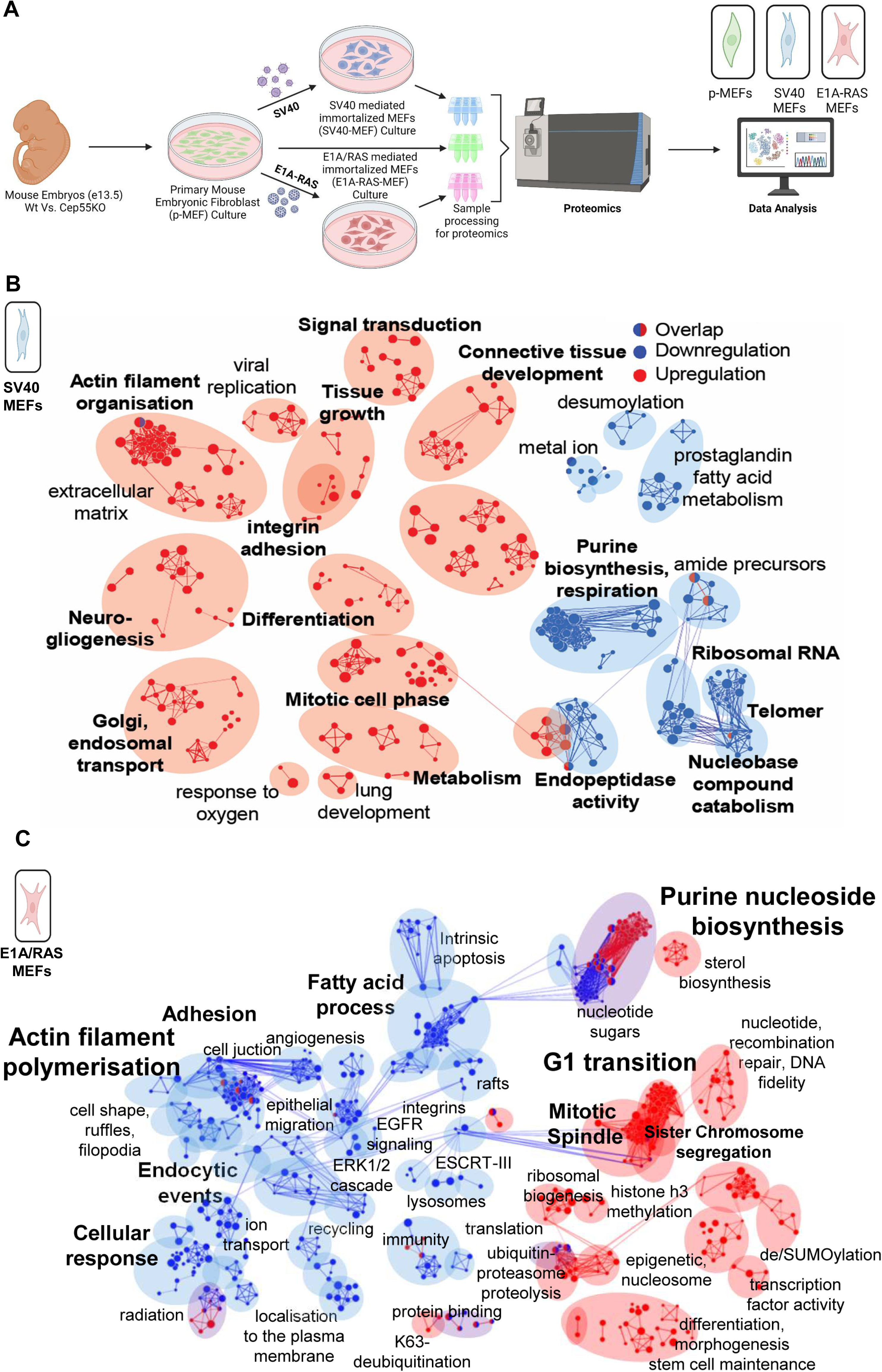

To investigate the biological context that exacerbate the oncogenic role of CEP55, we analyzed CEP55 correlation with 1,387 gene signatures representing oncogenic pathways and cancer hallmarks across 32 TCGA cancer types. These signatures encompassed critical processes such as cell cycle regulation, DNA replication/repair, immune signaling, apoptosis, oncogenic signaling, and metastasis-related processes. Spearman correlation analysis across RNA-seq datasets (n > 100 patients per cancer type) identified breast cancer, lung sarcoma, skin cutaneous melanoma, and thymic tumors the strongest correlations between CEP55 expression and oncogenic pathway activation (**Sup fig 1c, Sup material_2**). Functionally, CEP55, known as a centrosomal protein and master regulator of cytokinesis, was most strongly associated with pathways related to mitotic spindle formation, microtubule organization, and cell division (**Sup fig 1c**). Positively correlated pathways included cell cycle regulators (p53, RB, FOXM1), DNA damage response (ATM/ATR signaling, MRN complex), oncogenic signaling (MYC, PLK, Aurora kinases, p38 MAPK), immune pathways (IL-6, IL-2, TNF, MHC II), and apoptosis-related programs (**Sup Table 1**).

**Table 1.**
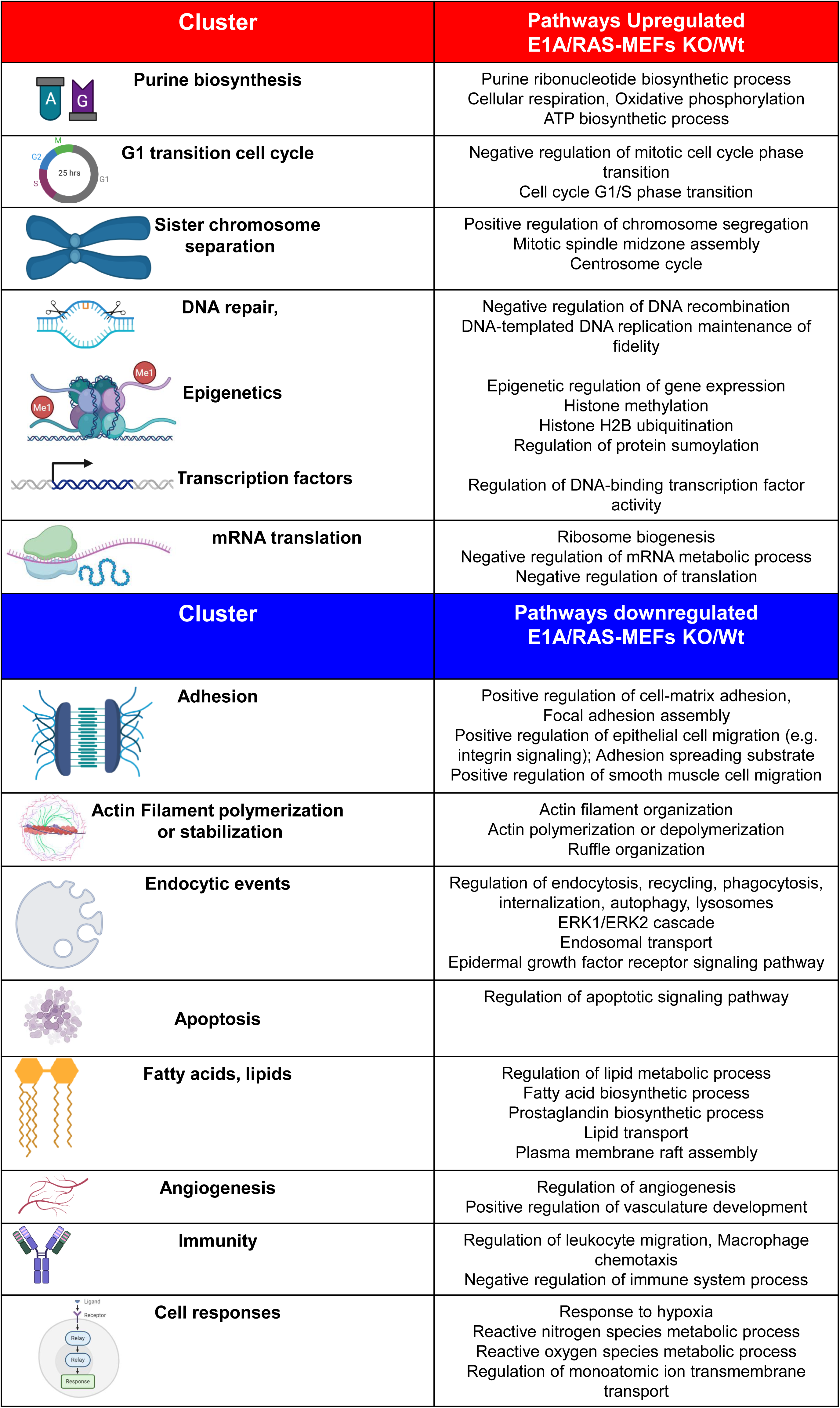
Differentially Up and Downregulated pathways in Cep55 E1A-Ras-MEFS.

### CEP55 expression is linked to immune cell infiltration and epigenetic modulation

We next examined CEP55 in the context of the tumor immune microenvironment (TIME), using data from CIBERSORT and TISIDB databases. High CEP55 expression positively correlated with the inferred abundance of several immune cell types, including activated memory CD4⁺ T cells, T follicular helper (T_fh) cells and Th2 cells (**Sup fig 1d, e**). Tumors with the highest immune correlation included cervical squamous cell carcinoma and endocervical adenocarcinoma (CESC), colon adenocarcinoma (COAD), head and neck squamous cell carcinoma (HNSC), kidney renal papillary cell carcinoma (KIRP), ovarian serous cystadenocarcinoma (OV) and thyroid carcinoma (THCA) (**Sup fig 1d, Sup material_3**). Analysis of CEP55 copy number variation and promoter methylation further revealed that CEP55 copy number gain was associated with modest immune cell changes, however, CEP55 promoter hypermethylation consistently correlated with reduced lymphocyte infiltration (**Sup material_1 Fig b**). This suggests that epigenetic silencing of CEP55 may suppress immune infiltration and contribute to immune evasion mechanisms in tumors.

CEP55 expression aligns with immune checkpoints and immune evasion signatures. Interestingly, CEP55 expression also correlated with the expression of multiple immune checkpoint molecules. Specifically, some of the tumors types with high CEP55 levels exhibited elevated expression of inhibitory checkpoint genes such as *TIGIT*, *LAG3*, *CTLA4*, *PD1* and, with a weaker but consistent co-occurrence with immunostimulatory ligands such as *CD80* and *CD86* (**Sup Fig 1e**). Notably, CEP55-high tumors displayed decreased *MHC-I* gene expression, suggesting impaired antigen presentation and a potential mechanism for immune evasion (**Sup material_1 Fig c**). Despite this reduction in *MHC-I*, expression of *TAP1* and *TAP2*, key genes in the antigen-processing machinery, remained positively correlated with CEP55, indicating that while antigen processing is retained, surface presentation may be limited. Furthermore, CEP55 expression demonstrated an inverse correlation with the immunosuppressive cytokine TGF-β1 and a positive correlation with pro-inflammatory chemokine CXCL8 (IL-8). In contrast, CCL14, a chemokine associated with monocyte recruitment, was lowest in CEP55-high tumors (**Sup material_1 Fig c-d**).

These findings support the existence of a complex immune contexture in CEP55-high tumors: while enriched for immune infiltrates, these tumors limit anti-tumor responses through immune checkpoints upregulation and reduced antigen visibility via low MHC-I. Furthermore, in thyroid, colon, ovarian, and cervical cancers, elevated CEP55 coincided with a Th2-skewed immune responses and was lowest in the C5 (immune-cold) subtype (**Sup material_1 Fig e**). In contrast in breast cancer, CEP55 was most highly expressed in the C2 subtype characterized by an IFN-γ–dominant immune response, and again lowest in the C5 subtype (**Sup material_1 Fig e**). These data suggest that CEP55 upregulation not only promotes proliferation and cell-cycle progression but also shapes an immunosuppressive tumor microenvironment linked to adverse clinical outcomes.

### Proteomic profiling of downstream effects upon Cep55 loss in untransformed and early-stage transformed mouse embryonic fibroblasts (MEFs)

To assess whether CEP55 ablation impact the cell proteome content and quality, we performed mass spectrometry-based proteomic analysis using primary or transformed MEFs, under Cep55 wildtype (Wt) or knockout (KO) conditions. Primary untransformed MEFs (pMEFs) were immortalised using the SV40 large T antigen, to mimic early transformation, or the adenoviral E1A/Ras^V12^ vector to mimic a tumorigenic stage, E1A/RAS (Rashidieh et al., 2021b). The workflow of proteomics and phosphoproteomics is illustrated in **Fig 1A**.

We first focused on the pMEF and SV40 conditions, identifying ∼2000 total proteins and 300-400 phosphoproteins. These analyses revealed significantly up- or downregulated proteins (q<0.05) upon Cep55 loss relative to Wt (**Sup fig 2a-b)**. The correlation between pMEFs-Wt and -KO proteome profiles (Spearman r ≈ 0.74) suggests that the lack of Cep55 causes protein expression changes, while normalized intensities across replicates indicate unchanged global proteome amount (**Sup fig 2c-d).** The cluster maps of enriched pathways in SV40-MEFs (early transformation stage, (**Fig 1B**) and E1A/RAS (tumorigenic stage, **Fig 1C**) suggested rewiring of several unexpected and separate biological processes in KOs.

**Figure 2:**
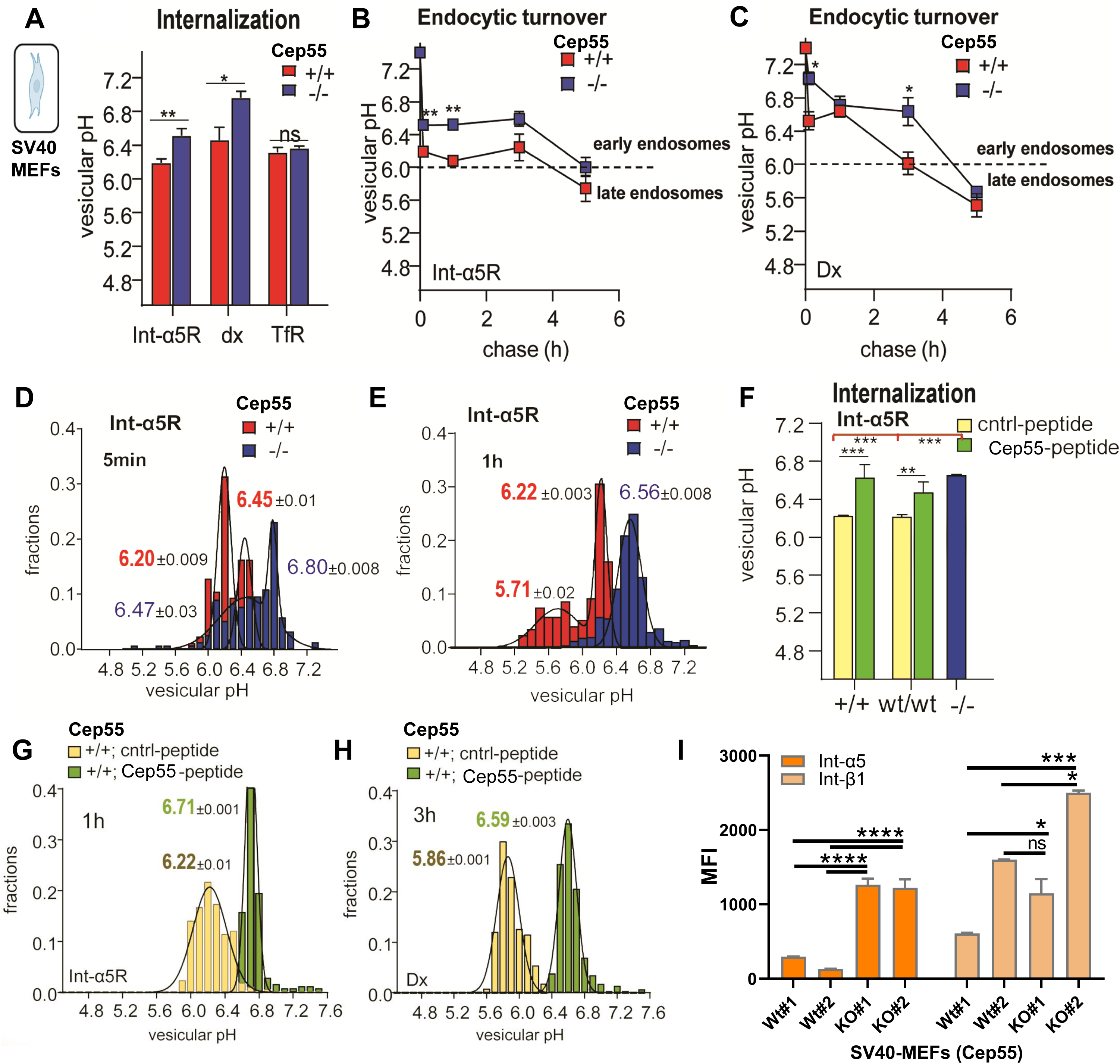

Downregulated processes in SV40-MEFs included ribosomal biogenesis, prostaglandin signalling, nucleotide metabolism, and fatty acid biosynthesis (blue, **Fig 1B).** Analysis of individual biological and molecular pathways in both pMEFs (untransformed) and SV40-MEFs revealed that significantly differentially downregulated proteins (log₂ fold change >L0.5, pL<L0.05) are involved in RNA processing, mRNA translation, and ribosomal biogenesis in Cep55-KO cells. Notably, ribosomal proteins such as Rpl24 and Rps23, along with several DEAD-box helicases implicated in ribosomal biogenesis and translational control, were significantly reduced in both KO conditions (**Sup fig 2 e-g; Sup fig 3a–d**). While some of these effects are likely indirect, they may also be linked to altered centrosome architecture. The centrosome and midbody have been shown to spatially localize specific RNAs (e.g. active translation of CHMP4B at midbody, a CEP55-linked protein), and to associate with spliceosomal components, often in conjunction with ribosomal proteins, ribonucleoprotein complexes, and elements of translational control (Zein-Sabatto and Lerit, 2021, Cerulo et al., 2023, Farmer et al., 2023).

Interestingly, SV40-MEF KO cells exhibited upregulation of pathways involved in actin filament organization, extracellular matrix (ECM) remodelling, integrin-mediated adhesion, connective tissue development, and neurogliogenesis (red, **Fig 1B**). Notably, the upregulated transforming growth factor beta (TGF-β) pathway driven by proteins such as cytokine TGF-β receptor II (Tgfbr2) and Cd109 is consistent with the observed inverse correlation between CEP55 and TGF-β expression (**Sup material_1 Fig c,d).** The enriched upregulated clusters in SV40-MEF KO (**Fig 1B**), and their pathways, summarized in **SupTable 2**, collectively suggest that Cep55 loss during early transformation modulates pathways involved in tissue growth, cell behaviour, differentiation, and immune regulation.

Remarkably, Cep55-KO in both pMEFs and SV40-MEFs resulted in significant upregulation of cell-substrate adhesion and ECM pathways (**Sup fig 3e-g**), in contrast to downregulation observed in tumorigenic E1A/RAS stage (**Fig 1C**). Key drivers of these processes included laminin and RGD-receptor integrins ItgaV, Itgb1 and especially in SV40-MEF-KO Itga5 (**Sup fig 3 e-f**), along with their ECM binding partner Thrombospondin 1 (Thbs1), and several collagens such as Col3a1 and Col1a1 (**Sup fig 3 g-h).** Upregulation of integrins (**Sup fig 3 e-f**) and collagens (**Sup fig 3 g-h)** suggested increased cell-cell and connective tissue interactions.

**Figure 3:**
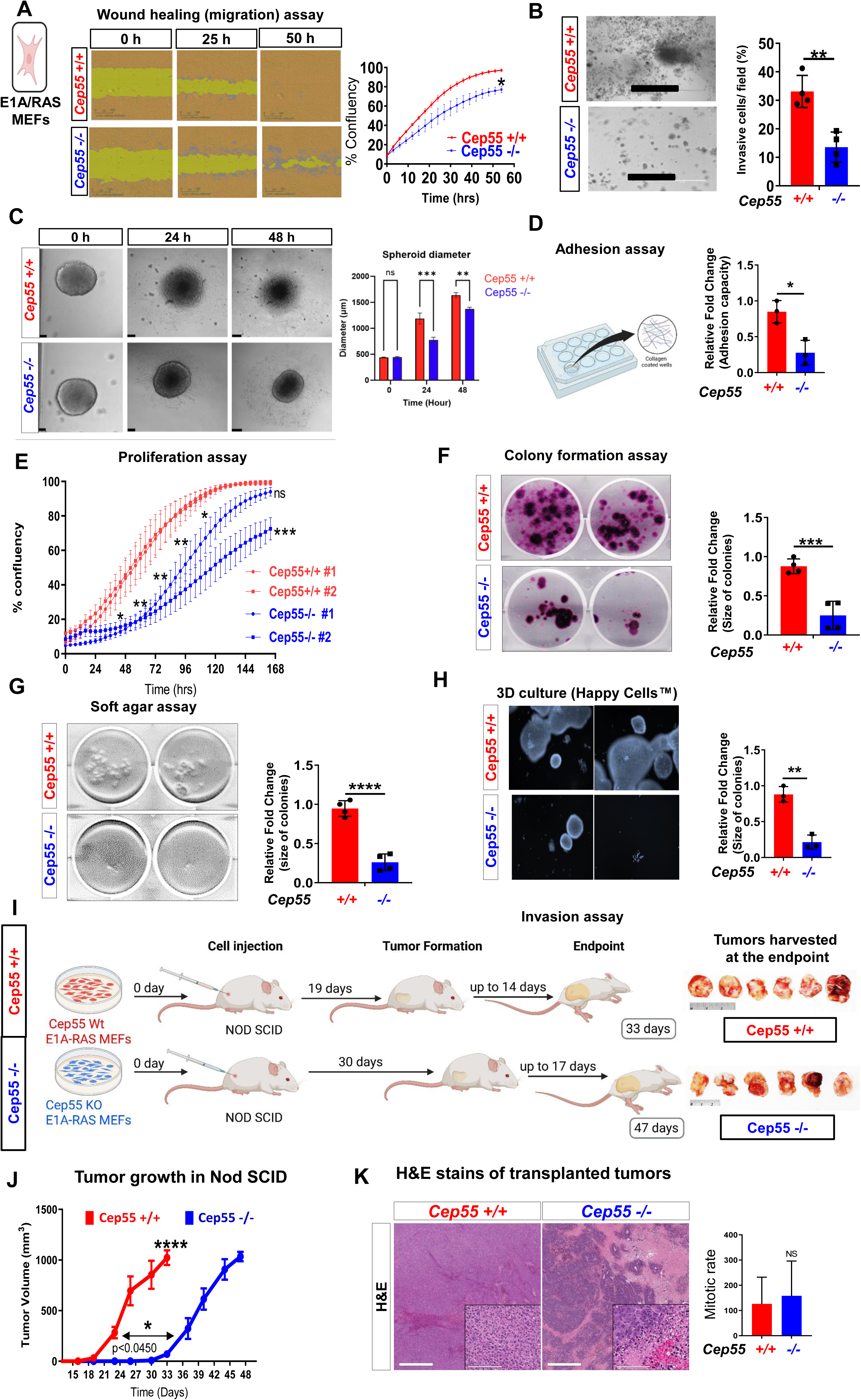

Both pMEF and SV40-MEF KO conditions also exhibited increased expression of variable actin-phospholipid membrane-binding proteins (**Sup fig 3 c-d**), again in contrast to tumorigenic E1A/RAS condition (**Fig 1C**). Notable examples include epidermal growth factor receptor (Egfr) and myristoylated alanine-rich C-kinase substrate (Marcks) in pMEF-KO, as well as Bridging Integrator 1 (Bin1) and cell adhesion molecule Kirrel1 in SV40-KO. Many of these proteins are either regulators or targets of endocytic processes. CEP55 directly interacts with and recruits the ESCRT-I (Endosomal Sorting Complex Required for Transport) component tumour-susceptibility gene 101 (TSG101) and ALG-2 interacting protein X (ALIX; PDCD6IP) to the midbody during cytokinesis (Lee et al., 2008, Elia et al., 2011). ALIX and ESCRT-I/II complexes cooperate to recruit the ESCRT-III subunit CHMP4B, facilitating abscission (Christ et al., 2016) Beyond their role in cytokinesis, ESCRT complexes are central to the endosomal sorting and degradation of ubiquitinated membrane proteins, including integrins, and are implicated in a wide range of cellular processes, from lipid, fatty acid, and amino acid metabolism to signaling, genome maintenance, and membrane repair (Apaja and Lukacs, 2014, Hurley et al., 2025). Proteomic heatmaps revealed reconfiguration of the endosomal and lysosomal adaptor and regulator proteins in Cep55-KO pMEF and SV40-MEF cells, including altered expression of ESCRT-0 to -III subunits such as Stam2 and Hrs (Hgs) (**Sup fig. 4a–c**). Notably, Chmp4b (ESCRT-III) was downregulated, and regulators of integrin trafficking, such as Ptpn23 (Hd-ptp) (Kharitidi et al., 2015), showed differential expression. In summary, proteomic profiling of Cep55-KO primary (pMEFs) and early-transformed (SV40-MEFs) suggests that reprogramming of the endosomal pathway may underlie the elevated expression of certain signalling and adhesion receptors (e.g. integrins) and contribute to the observed metabolic, developmental and differentiation-related processes (**Sup fig 2e-g**).

### CEP55 loss drives distinct proteomic and phosphoproteomic changes in tumorigenic E1A/RAS MEFs

We next focused on the impact of CEP55 loss at the tumorigenic stage using E1A/Ras-transformed MEFs, which model oncogenic transformation. Following the workflow in **Fig 1A**, we analyzed ∼5,000 total proteins and phosphopeptides with ∼500 phosphosites. Compared to primary (pMEFs) and early-transformed (SV40-MEFs) (**Fig 1B, Sup fig 2e-f and 3a-d**), E1A/Ras-transformed MEFs exhibited substantial differences in enriched clusters and biological and molecular pathways (**Fig. 1C, Sup fig 2g and 5 a-d**). These upregulated and downregulated pathways and clusters are summarized in Table 1. Strikingly, Cep55-E1A/Ras KO cells exhibited downregulation of pathways related to cell adhesion, endocytosis and oxidative stress response, which contrasted with upregulation of these pathways observed in pMEF and SV40-KO cells (**Fig 1C, Sup fig 5a-b)**. Processes such as integrin-mediated signalling, focal adhesion and integrin binding were decreased, driven by proteins such as integrins ItgaV, Itgb3, Itgb5, and focal adhesion protein tyrosine kinase 2 (Fak; Ptk2) and late-endosomal tetraspanin Cd63 (**Sup fig 2g**, **Sup fig 5c-d**). While ECM binding components were decreased, the amount of collagen remained unchanged in E1A/Ras KO (**Sup fig 5e).**

In contrast, the upregulated pathways in E1A/Ras-CEP55 KO cells involved chromatin remodelling, nucleosomal DNA binding (e.g., histones h1-0, h1-1), and regulatory proteins including histone methyltransferase SET Domain Containing 2 (Setd2), serine/threonine-protein kinase tousled-like 2 (Tlk2), and HAUS-family spindle assembly proteins. These findings suggest a shift in epigenetic regulation, DNA damage response, and mitotic control (**Sup fig 2g and 5a-b)**. Phosphoproteomic analysis further revealed increased phosphorylation of proteins involved in chromatin dynamics, cytokinesis, and mRNA processing, while phosphorylation was decreased in pathways related to actin organization, transcription elongation, and translation regulation (**Sup fig 5c-d**). As expected, the downregulation of Akt1 phosphorylation (Ser^473^) and mitosis-specific Cep55 (Ser^426^) phosphorylation was observed. Phosphorylation was notably reduced in several sites of Bridging integrator 1 (Bin1), a tumour suppressor that binds Myc, and Fibroblast growth factor receptor substrate 2 (Frs2), a key signal-transducing adaptor protein involved in the MAPK/ERK, PI3K/AKT/mTOR signalling (Zhou et al., 2009). Specific downregulated Frs2 phosphosites included (T132; T135; T137, S211, S365) along with phosphosites of cytoskeletal protein Pdlim2 and vesicular trafficking and cilium assembly protein Rab3ip (**Sup fig 5e**, **Sup Table 3**). In contrast, increased phosphorylation was observed in the Breakpoint cluster region kinase (Bcr; at multiple sites including S461 and S464) and DNA methyltransferase 1 (Dnmt1) in E1A/Ras KO MEFs. Other proteins with significantly upregulated phosphorylation included microtubule-associated proteins (Kif23), centrosome-endosomal proteins (Cc2d1a) and signalling proteins (Pak1) (**Sup fig 5e**, **Sup Table 3**). Notably, some phosphosites involved in chromatin binding (Hmga1), DNA repair (Tnks1bp1), mRNA splicing (Srrm1 and 2), stress-responsive translation (Ubap2l), cell shape and motility (Myh9, Map7d1) were consistently up- or downregulated across pMEF, SV40, and E1A/Ras KO MEFs (**Sup fig 5h**). Phospholipase A2 IVA (Pla2g4a; cPLA2α) was downregulated at both total protein and phosphorylation in all KO conditions (**Sup fig 5h**) correlating with suppression of the prostaglandin/fatty acid metabolism pathway (**Sup fig 2g**).

Together, these data demonstrate that genetic ablation of Cep55 in an oncogenic background leads to distinct proteomics and phosphoproteomic alterations in comparison to primary and early transformed MEFs. These changes likely reflect the altered signalling and metabolic landscape characteristic of fully transformed cells.

### CEP55 expression modulates integrin receptor endocytosis, signalling and cell migration

To functionally validate Cep55 KO-associated alterations in early transformed (SV40) MEFs, we analyzed the effect of Cep55 loss on integrin receptors, a connection not yet expolored. Several RGD-motif binding integrin receptors, such as integrin α5β1, undergo enhanced internalization and recycling in oncogenic cells, a process linked to increased cell migration, cancer progression, and metastasis. Tight regulation of dynamics in endocytosis, actin cytoskeleton, ligand binding, ECM interaction and integrin clustering is essential to coordinate integrin activation (Chastney et al., 2025, Kharitidi et al., 2015).

To assess integrin trafficking, we leveraged the pH-dependent nature of different endosomal compartments to quantitatively track cargo at the single-vesicle and single cell level using pH-sensitive, fluorescence-conjugated probes combined with fluorescence microscopy (Barriere and Lukacs, 2008, Kharitidi et al., 2015, Kazan et al., 2019). Transferrin receptor (TfR) served as a control for fast, ubiquitin-independent endocytosis and dextran (Dx) for non-specific endocytosis. Labelled integrin α5β1 (IntαR) endocytosis in early transformed SV40 Wt (+/+) MEFs revealed that internalized integrins localize to compartments with overlapping recycling (∼pHL6.3) and early endosomal (pHL6.0–6.4) pH profiles, consistent with signalling-competent integrin trafficking dynamics (**Fig 2A-B**).

Cep55 loss significantly delayed integrin α5β1 internalization and its targeting to early and late endosomes (pHL<L6) (**Fig 2A-B**). In Cep55 KO (−/−) cells, integrin α5β1 exhibited prolonged residence in sub-plasma membrane vesicles (pHL>L6.5), whereas in Wt (+/+) cells, the receptor localized to recycling compartments (pHL6.3) and late endosomes (pHL5.1) (**Fig 2D-E**). Immunofluorescence microscopy of internalized integrin α5β1 receptor confirmed its colocalization with the early endosomal marker EEA1 in Wt cells, but not in Cep55 KO cells (**Sup fig 6a**). Furthermore, dextran internalization and endocytosis were delayed in Cep55 KO (−/−) cells (**Fig 2A, C**), suggesting that Cep55 alters endosomal organelle structure and dynamics.

To test whether disrupting the interaction between Cep55 and the ESCRT adaptor Alix (PDCD6IP), critical for abscission, affects integrin trafficking, we used an ALIX-peptide (GPPSLNY) (Lee et al., 2008) that blocks the Cep55–Alix binding site interaction and a control peptide with alanine substitution (GPASLNA) in two SV40-transformed wild-type (Wt/Wt, +/+) cell lines. Endocytic pH analysis revealed that this Cep55-peptide mimicked the KO phenotype, significantly delaying integrin α5β1 trafficking and resulting in sub-plasma membrane vesicle localization (pH 6.6–6.7) (**Fig 2F–G**), also delaying Dx marker to recycling compartment (∼pH 6.3) after 3 h instead of late endosomes (**Fig 2H**). FACS analysis confirmed increased cell surface expression of integrin α5 and β1 in SV40 Cep55-KO MEFs, consistent with proteomics findings (**Fig 2I, Sup fig 6b**).

Since integrin endocytosis was impaired in SV40 KO cells, suggesting defective integrin activation, we next assessed downstream signalling. In SV40 KO cells, phosphorylated Akt (Ser^473^) and phosphorylated ERK1/2 (Phospho-p44/42 MAPK, Thr^202^/Tyr^204^), both key β1-integrin signaling effectors (Velling et al., 2004), were downregulated in a Cep55 expression dependant manner while the total proteins and upstream mediators remained unchanged (**Sup fig 6c**). Similar trends were evident in E1A/RAS-transformed MEFs, where despite comparable KRAS expression across conditions, Cep55 KO led to decreased levels of phosphorylated ERK1/2, integrin β5, phosphorylated focal adhesion kinase (pFAK=p-Ptk2),, phosphorylated Paxillin, and RHOA (**Sup fig 6d**). Notably, in E1A/RAS MEFs, Cep55 deletion also reduced expression of integrin α5, a subunit frequently upregulated in oncogenic contexts. These findings indicate that Cep55 is critical for maintaining integrin activation and downstream signaling required for oncogenic signaling in transformed cells.

To examine the functional consequences of Cep55 loss on cell motility, we performed time-lapse single-cell tracking using the holomonitor live-cell imaging. Cep55-KO cells exhibited significantly reduced motility, with approximately 20–30% lower average migration speed and total distance travelled per cell compared to wild-type controls (**Sup fig 6e**). These findings indicate that Cep55 loss primarily impairs migration velocity, likely as a consequence of disrupted integrin trafficking and signaling. Taken together with our previous observations, the data show that Cep55-KO disrupts integrin endocytosis, recycling, and activation, leading to attenuated focal adhesion signalling as evidenced by decreased phosphorylation of FAK and paxillin. Concurrent suppression of RHOA and pERK1/2 signalling, likely contributes to altered cytoskeletal organization and reduced migratory capacity. These data support a model in which Cep55 is essential for sustaining integrin-dependent adhesion and signaling required for efficient cell migration.

To explore the clinical relevance of these findings, we queried the cBioPortal and The Cancer Genome Atlas (TCGA) datasets to assess whether CEP55 expression correlates with integrin signalling components in human cancers, particularly in breast cancer, where integrin signalling is a well-established driver of disease progression. We analyzed a cohort of 963 patients with invasive breast carcinoma, focusing on mRNA expression and copy number alterations for genes including CEP55, integrins αV, α5, β1, fibronectin (FN1), and FAK (PTK2). Our analysis revealed strong co-occurrence and positive correlation between CEP55 expression and the expression of integrins β1, αV, α5, fibronectin, and FAK (**Sup fig 6f; Sup Table 4**). These results suggest that the relationship between CEP55 and integrin-mediated signaling observed in our experimental models may be conserved in human cancers, supporting a broader role for CEP55 in regulating adhesion and signaling networks that drive tumor progression.

### Cep55 Knockout impairs cell migration, transformation and invasive growth *In Vitro*

To evaluate the role of Cep55 in integrin-regulated cellular behaviours, we assessed motility, migration, and invasion in oncogenic E1A/Ras MEFs. In wound-healing assays, Cep55-KO (−/−) cells exhibited significantly impaired migration, closing approximately 40% less wound area than Wt (+/+) cells after 50 hours (**Fig 3A**). In our custom 3D spheroid invasion assays using PIC-GFOGER hydrogel matrices, Cep55-KO cells showed reduced invasion areas at 24 and 48 hours post-embedding compared to oncogenic Wt cells (**Fig 3B**). We examined several hydrogel conditions for the 3D spheroid invasion assay and tailored the hydrogel physicochemical properties and microenvironment to support optimal spheroid invasion conditions. We found that the combined matrix stiffness of 330 Pa, 1% GFOGER ligand density and strain stiffening onset of 37 Pa provided a scaffold that supported spheroid stability, growth, and cell migration through the hydrogel microenvironment. (**Fig 3C**). Moreover, a collagen adhesion assay revealed significantly reduced adhesion of KO MEFs (**Fig 3 D**). Collectively, these findings confirm that Cep55 supports integrin-driven migration, adhesion and invasive capacity in oncogenic MEFs.

To assess whether Cep55 loss also affects tumorigenic characteristics, we next analyzed proliferative capacity and clonogenicity potential in E1A/RAS MEFs, Real-time live-cell imaging (IncuCyte®) revealed that Cep55-KO^−/−^ MEFs proliferated at ∼50% the rate of Cep55-Wt^+/+^ cells, with noticeable growth defects emerging by 48–72 hours and a ∼2-fold reduction in confluence observed by days 5–7 (p<0.001) (**Fig 3E**). Clonogenicity was also significantly compromised in Cep55-KO (−/−) MEFs compared to their Wt (+/+) counterparts, as evidenced by diminished colony formation in both 2D and 3D assays (**Fig 3F).** In soft agar, Cep55-KO MEFs formed ∼70% fewer colonies, which were markedly smaller, typically less than one-third the size of Wt colonies (**Fig 3G**). To further assess 3D growth in a suspension environment, we used Happy Cell Advanced Suspension Medium® (HCM), a low viscosity liquid reagent matrix that enables spheroid formation from single cells (Takahashi et al., 2018). Similarly, in Happy Cell matrix, Cep55 KO E1A/Ras MEFs yielded dramatically fewer and smaller spheroids compared to Wt (**Fig 3H**). These results indicate that Cep55 is essential for anchorage-independent growth and survival under non-adherent conditions, underscoring its role in supporting anoikis resistance and tumorigenic potential.

### Loss of CEP55 Delays tumour growth and progression *In Vivo* (Cell Transplant Model)

Our *in vitro* findings demonstrate that Cep55 deficiency impairs multiple oncogenic behaviors in MEFs, including integrin-mediated AKT and FAK signaling, adhesion, invasion, and anchorage-independent growth. To determine whether these impairments translate to reduced tumorigenicity *in vivo*, we employed a cell transplant model. E1A/Ras-transformed MEFs (either Cep55-Wt (+/+) or Cep55-KO (−/−) were injected intraperitoneally (IP) into 8–10-week-old immunodeficient NOD/Scid mice (nL=L6 per group All mice injected with Cep55 Wt (+/+) MEFs developed palpable tumors by ∼19 days post-injection. In contrast, mice injected with Cep55 KO (−/−) MEF developed palpable tumors only after ∼30 days (p<0.04), a greater than 50% delay in tumor onset (**Fig 3I**). Moreover, once tumors were established, those derived from Cep55-KO cells grew more slowly, reaching the experimental endpoint (1000Lmm³) by day 47 on average, compared to day 33 for Wt tumors (pL<L0.01). This delay in tumor progression was reflected in overall tumor growth kinetics (**Fig 3I**).

Histological examination, H&E-stained sections, of endpoint tumors revealed large necrotic regions in Cep55-KO tumors, whereas WT tumors were more uniformly viable (**Fig 3K, right panel**). Interestingly, the mitotic index (mitoses per high-power field) was not significantly different between WT and KO tumors (**Fig 3K, left panel**), suggesting that proliferation of the viable tumor cells *in vivo* was similar at endpoint (tumors harvested at different endpoints). However, the greater necrosis in KO tumors may reflect more cell death and possibly due to impaired vascularization or nutrient supply.

Together, these data demonstrate that Cep55 loss significantly delays tumor initiation and slows tumor progression in a transplant setting, consistent with the *in vitro* findings of impaired growth and survival. Combined with our unbiased proteomic data, and bioinformatic analyses of human cancer databases, these findings highlight a critical and previously underappreciated role for Cep55 in regulating cell adhesion and integrin signalling via the endocytic and ESCRTs pathway. Loss of Cep55 impairs integrin activation and downstream oncogenic signaling, culminating in suppressed tumor growth and invasive capacity. These results suggest that CEP55 may represent a novel target for therapeutic intervention in integrin-driven malignancies.

### A Novel Mouse Model of Inducible Cep55 cKO Marked Cep55 as Dispensable for Normal Adult Viability

Building on the anti-tumor effect of Cep55 loss in our transplant model, we next employed genetically engineered mouse model (GEMM) to further investigate its role in tumorigenesis *in vivo*. Our previously reported constitutive Cep55 knockout mouse was perinatal lethal, largely due to neurodevelopmental defects during embryogenesis, consistent with the known essential role of Cep55 during early development (Rashidieh et al., 2021b). To overcome this limitation, we generated an inducible *Cep55* conditional knockout (cKO) in adult mouse using a *knockout-first* allele amenable to the Cre-LoxP system (**Sup fig 7a**).Through a breeding strategy (**Sup fig 7b**), we crossed *Cep55 ^Fl/Fl^* mice with Rosa26-CreERT2 mice to obtain *Rosa26-CreERT2^+;^ Cep55^Fl/Fl^* animals. In this system, Cre recombinase is expressed ubiquitously but remains inactive until tamoxifen administration, enabling controlled *Cep55* deletion in adult tissues. To assess *Cep55* expression, we first analyzed publicly available murine RNA-Seq Cap Analysis of Gene Expression (CAGE) datasets which revealed high *Cep55* RNA expression levels in testis and moderate levels in spleen, lung and intestine in adult mice (**Sup fig 7c**). We confirmed these findings at the protein level using immunobloting in 8-week old mice. The highest expression of Cep55 was observed in testis, with moderate expression in the spleen and intestines, and undetectable levels in the other tissues analyzed (**Sup fig 7d**). Efficient CEP55 protein depletion following tamoxifen administration was confirmed in the spleen (**Sup fig 7e**). To further validate the model *in vitro*, we isolated MEFs from *Rosa26-CreERT2^+;^ Cep55^Fl/Fl^* embryos and induced knockout using 1 µg 4-hydroxytamoxifen (4OHT) for 72 hours. This induced robust Cep55 loss and significantly impaired cell proliferation, replicating the phenotype seen in constitutive Cep55 knockout MEFs (**Sup. Fig. 6f**).

Next, we evaluated the physiological impact of Cep55 deletion *in vivo* in adult mice. Eight-week-old Rosa26-Cre^ERT2^; Cep55^Fl/Fl^ mice were treated with tamoxifen (1mg) to induce global Cep55 knockout (n=8), alongside littermate controls (Cre+ PBS injected, or Cre-Cep55^Fl/Fl^ tamoxifen-treated; n=6). Remarkably, all Cep55-cKO mice remained viable, healthy (weight and appearance) and behaviourally normal for at least 25 weeks post-induction, with no overt phenotype. At necropsy, major organs appeared normal, with no gross morphological defects after Cep55 deletion (**Sup fig 7f**). These results indicate that Cep55 is dispensable for adult tissue homeostasis, consistent with its low baseline expression in most adult tissues except testis. This supports the notion that anti-tumor effects of Cep55 loss are tumor-specific and highlights CEP55 as a promising therapeutic target with minimal expected toxicity in normal adult tissues.

### Genetic ablation of Cep55 prolongs survival in a Pten-deficient tumor-prone mouse model

Previous studies have shown that CEP55 promotes PI3K stabilization and enhances AKT activation (Chen et al., 2007,Sinha et al 2020) functionally mimicking the pathway activation caused by PTEN loss. In this context, CEP55 may become particularly critical to maintain hyperactivated PI3K/AKT signaling and promote malignancies. We therefore hypothesized that genetic ablation of *Cep55* in a *Pten-KD* background could suppress tumor progression by blunting PI3K/AKT, offering an alternative strategy where direct inhibitors of this pathway have shown limited clinical success.

To test this, we generated a conditional, tamoxifen-inducible mouse model allowing the co-deletion of Cep55 and Pten in adult tissues enabling us to examine whether inducible *Cep55* KO could reduce tumor incidence or delay onset in a PTEN-deficient tumor prone model (**Fig 4A**). We crossed our Cep55 ^Fl/Fl^ line with Pten^Fl/Fl^ mice to allow simultaneous deletion of both genes. Because Cep55 (locus 19C2) and Pten (locus 19C1) reside on the same chromosome, we calculated a ∼3% likelihood of recombination to obtain the floxed alleles in cis. Out of ∼200 pups genotyped, we identified one with a successfully recombined Cep55-Pten^Fl/+^ allele, which was subsequently bred with RosaCreERT2 mice. This breeding strategy produced RosaCreERT2*^+/−^*-Cep55 *^Fl/Fl^*-Pten^Fl/Fl^ offspring (hereafter referred to as Cep55-Pten: CP-DKD) and *Pten^Fl/Fl^*-*RosaCre^ERT2+/−^*. (Pten KD). Following tamoxifen administration at 6 weeks of age, we confirmed efficient deletion of both genes by immunoblotting across some tissues with Cep55 visible expression including spleen, liver, and testis (**Sup fig 8a**).

**Figure 4:**
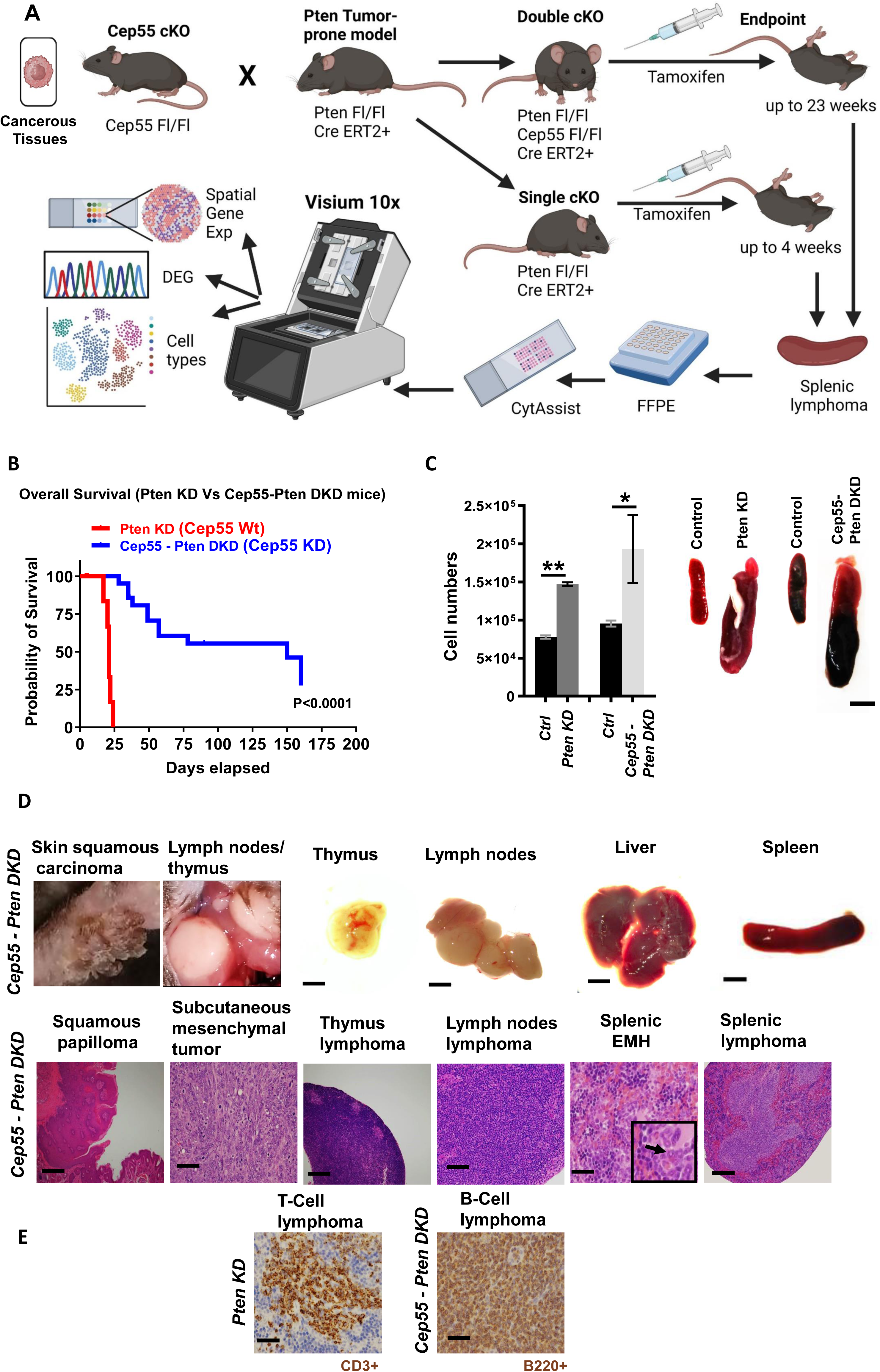

Consistent with previous reports, tamoxifen-induced (Cre ERT2 recombination) Pten-KD mice developed aggressive lymphomas exhibited a median survival of only 21 days (Backman et al., 2001; Suzuki et al., 2001). In contrast, Cep55-Pten DKD mice showed strikingly prolonged survival, with a median of ∼150 days, a greater than 7-fold extension (p < 0.0001, log-rank test; **Fig 4B**). While all Pten KD mice died within four weeks post-induction, approximately 20% of CP-DKD mice remained alive and lymphoma-free at the 160-day experimental endpoint. This dramatic survival benefit indicates that loss of Cep55 greatly delays and hamper the tumorigenesis driven by complete Pten loss.

Pathological examination of moribund mice confirmed extensive splenomegaly and T-cell lymphomas in Pten KD animals (**Fig 4C**). In contrast, CP-DKD mice displayed more heterogeneous and delayed-onset pathologies. While thymic and splenic lymphomas were still observed in some cases, many CP-DKD mice developed benign or low-grade lesions in other tissues at later time points including uterine endometrial neoplasms (females), skin papillomas, (by 4-5 months) and rare cases of prostatic hyperplasia (males) (**Fig 4D–E; Sup Table 5**). Notably, female CP DKD mice exhibited earlier mortality, with 100% dying by 7 weeks (often with T-cell lymphomas), whereas 60% of male CP DKD mice survived beyond 5 months, suggesting sex-specific influences on tumor development. These slower, often benign lesions suggest that in the absence of Cep55, the rapid lethal lymphomagenesis driven by Pten loss is mitigated, unmasking later-onset pathologies in other organs. Surviving CP DKD mice (especially males) displayed unusual phenotypes mentioned above including skin and prostate lesions, hinting that CEP55 may also impact specific tissue contexts differently (**Fig 4D–E**; **Sup Table 5**). Collectively, the data suggest *Cep55* can delay tumorigenesis and tumor latency on a *Pten*-deficient background.

At a molecular level, Cep55 loss dampened oncogenic signaling in Pten-KD tumors. Immunohistochemistry (IHC) of spleen and liver tissues showed that CP-DKD tumors had significantly reduced levels of phosphorylated AKT (Ser^473^), phospho-ERK, and β-catenin compared to Pten KD tumors (**Sup fig 8b-c**). FAK expression was also reduced in a subset of CP-DKD tumors (**Sup fig 8d**), supporting the role of CEP55 in amplifying PI3K/AKT, MAPK, and focal adhesion signaling *in vivo*.

Together, these findings demonstrate that CEP55 is a critical enhancer of PI3K/AKT-driven tumorigenesis in the context of PTEN loss. Its deletion significantly prolongs survival and alters tumor progression, positioning CEP55 as a viable therapeutic target in PI3K pathway– driven malignancies.

### Spatial Transcriptomics Reveals Immunoregulatory and Differentiation Programs in CP-DKD Tumors

To further investigate the molecular basis of delayed tumorigenesis in CP-DKD mice, we performed spatial transcriptomic profiling using CytAssist2 and Visium platforms following the described workflow (**Fig 4A**). Data were normalized using nCount and nFeature metrics to control for sequencing depth and gene complexity, with Hb% included to account for hemoglobin transcript abundance minimizing bias from highly expressed erythrocyte-derived genes (**Sup fig 9a**). Compared to PTEN-KD tumors, CP-DKD tumors exhibited significant upregulation of genes (Average log2 fold change > 2.5) associated with immune regulation (e.g., *Cd55*, *Igha*, *Dapl1*) and downregulation (Average log2 fold change −2.5) of cell cycle, cytokinesis, and ECM-related genes (e.g., *Col1a1*, *Col1a2*, *Clec9a, Ptp1*) (**Fig 5A–C**). Notably, *Aida* (JUN/JNK signaling), *Cd55* (immune evasion), and *Dapl1* (epithelial differentiation) were among the most upregulated genes. Downregulated genes included collagen family members, *CD207* (T-cell activation), and *SERPINE1* (glioma progression), while cancer/testis antigens *EIF2S3Y*, *DDX3Y*, and *KDM5D* were also differentially expressed (**Fig 5A-C, Sup material_4**).

**Figure 5:**
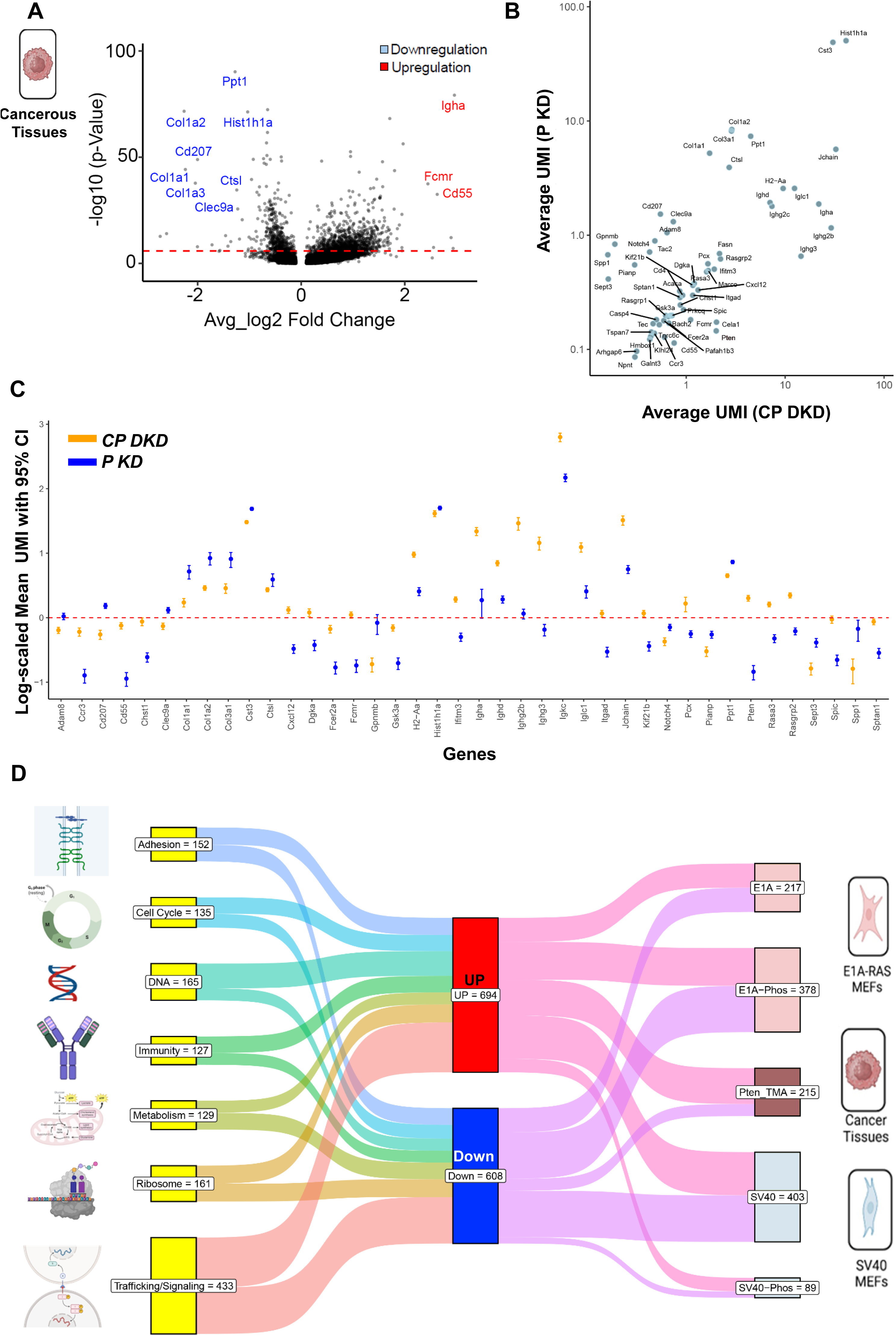

Within the annotated splenic lymphoma regions, *Cep55* transcripts levels were significantly reduced in Cep55-Pten DKD tumors compared to Pten-KD tumors, with partial re-expression observed in some tumors (**Sup fig 9b**). These transcript-level findings were validated by IHC for CEP55 protein (**Sup fig 9c**). Pten transcript was low in Cep55-Pten DKD tumors but protein levels were undetectable or nonspecific (**Sup fig 9d,e**). B cells enrichment in CP-DKD tumors was evidenced by upregulation of protein tyrosine phosphatase receptor type C gene (*Ptprc)* and confirmed by increased B220 protein levels on IHC (**Sup fig 9f**). Ptprc (CD45), is a transmembrane phosphatase expressed across hematopoietic cells, including B cells, T cells, and other leukocytes, is critical for immune signaling (Rheinländer et al., 2018). Interestingly, the B220 isoform of Ptprc is commonly used to identify B cells in mice, linking Ptprc directly to B cell profiling in both transcriptomic and protein-level analyses.

Pathway analysis revealed upregulation of CD4+ T-cell differentiation, integrin-mediated cell adhesion, myeloid cell development, platelet activation, TORC1 signalling, and the p38 MAPK cascade in PTEN-KD tumors relative to CP-DKD tumors. In contrast, pathways related to cytokinesis, microtubule cytoskeleton organization, and cell cycle checkpoint signaling were downregulated (**Sup fig 10a-c**). Corresponding changes in molecular functions, including those involving integrin, fibronectin and growth factor binding, Rac1, RhoA, Fgfr1 and death receptor signaling, were also observed (**Sup fig 10a-e**).

Comparison of unique differentially expressed genes in stroma and tumor regions highlighted changes in tumorigenesis-related genes, such as downregulation of *MMP2*, *MMP12*, *Spp1*, and upregulation of *Gpd1*, *Cdo1*, and *Il7r* (**Sup fig 10d**). The dual roles of these genes in tumor progression underscore the importance of evaluating transcriptomic signatures within cellular context.

Strikingly, key pathways disrupted in CP-DKD tumors including integrin-mediated adhesion, cytokinesis, and cell–substrate interactions were also dysregulated in our proteomic analysis of Cep55-deficient E1A/RAS MEFs (**Sup Table 6**). These findings were validated against TCGA data, confirming consistency across mouse and human cancer studies (**Fig 5D; Sup material_ 5**).

In summary, spatial transcriptomics revealed that Cep55 loss in a PTEN-deficient background induces a profound shift in the tumor microenvironment, characterized by immune activation, differentiation programs, and suppression of proliferative and invasive signaling. These findings highlight CEP55 as a crucial amplifier of PTEN-deficient tumorigenesis and a potential therapeutic target, particularly for cancers resistant to conventional PI3K/AKT-directed therapies.

## Discussion

CEP55 overexpression has been implicated in tumorigenesis across a broad range of cancers, including T-cell lymphoma, breast, lung, colorectal, and liver cancers. Traditionally recognized as a cytokinesis regulator in cancer cells; CEP55 is emerging as a multifunctional oncogenic driver with expanding roles in tumorigenesis (Sinha et al., 2020). It promotes malignant transformation by activating prosurvival pathways such as PI3K/AKT and contributing to genomic instability.

Here, we provide comprehensive *in vitro* and *in vivo* evidence that, while Cep55 is essential during embryonic development, it is largely dispensable for normal adult tissue homeostasis Using an inducible Cep55 cKO mouse model, we circumvented the limitations of perinatal lethality observed in constitutive knockouts and showed that its loss in adult mice had no overt physiological impact (Rashidieh et al., 2021b, Wangmo et al., 2024, Tedeschi et al., 2020). This finding aligns with expression analyses showing high Cep55 expression confined to testis, with moderate levels in the adult spleen and intestine, supporting a limited role in normal adult physiology.

Proteomic profiling of Cep55-deficient mouse embryonic fibroblasts (MEFs) revealed transformation stage-specific alterations in key cellular pathways. In primary and early-transformed MEFs, Cep55 loss led to downregulated ribosome biogenesis and RNA processing pathways, implicating a role in translational control and cellular metabolism. These effects may be mechanistically linked to CEP55’s centrosomal and midbody functions. These structures are known to spatially organize RNA and protein complexes involved in cell division and growth (Zein-Sabatto and Lerit, 2021, Cerulo et al., 2023, Farmer et al., 2023).

Interestingly, in early-transformed cells, Cep55-deficiency led to upregulation of extracellular matrix (ECM) remodeling, integrin signaling, and actin cytoskeleton organization. However, these pathways were suppressed in fully transformed, tumorigenic MEFs, while chromatin remodeling and DNA damage response pathways were enriched. This shift likely reflects the complex interplay between oncogenic signaling and epigenetic regulation during advanced tumorigenesis. Functional analyses revealed that Cep55 regulates integrin α5β1 trafficking through the endosomal pathway, in-part via its interaction with ESCRT proteins (ALIX and TSG101) (Lee et al., 2008). Loss of Cep55 impaired integrin internalization, prolonged retention near the plasma membrane, and defective trafficking to early and late endosomes, paralleling delayed endocytosis of other cargos such as dextran. These findings support a model where Cep55 coordinates membrane remodeling and cargo sorting at endosomal compartments, consistent with known roles of ESCRT complexes beyond cytokinesis (Lee et al., 2008, Christ et al., 2016, Apaja and Lukacs, 2014).

Importantly, disruption of the Cep55-ALIX interaction phenocopied impaired integrin internalization seen in cells with Cep55-loss, underscoring the functional importance of this axis. Notably, integrin signaling has been linked to centrosome integrity (Combedazou et al., 2020) and separation during early mitosis in human cells (Kamranvar et al., 2022) processes that may be compromised in the absence of CEP55. Supporting this notion, Kamranvar et al. (2016) reported that inhibition of integrin/FAK signaling delays abscission by affecting CEP55 recruitment to the midbody. Our findings suggest a reversal of this relationship: in the absence of Cep55, integrin–FAK signaling is impaired in transformed-MEFs. This points to a potential feedback loop wherein Cep55 promotes integrin signaling, and active integrin pathways may, in turn, support Cep55 function during cell division. Consistent with defective integrin trafficking, *Cep55*-deficient MEFs exhibited reduced activation of downstream signaling components, including diminished phosphorylation of Akt, Erk, FAK, and Paxillin. These molecular impairments translated into reduced cell migration, adhesion, invasion, and clonogenic growth in both 2D and 3D culture assays.

Corroborating the *in vitro* findings, Cep55-KO MEFs (E1A/RAS transformed) formed tumors with delayed onset, slower growth, and increased necrosis, suggesting compromised tumor fitness. Notably, the mitotic index remained unchanged at endpoint, indicating that reduced tumor progression is likely driven by impaired survival, adhesion, or microenvironment interactions rather than proliferation alone. These data highlight the importance of Cep55 in supporting integrin-mediated processes critical for tumor progression. These experimental observations were reinforced by bioinformatic analyses, which revealed co-occurrence and positive correlation between CEP55 copy number gains and integrin signaling components in human breast cancer datasets. This supports the hypothesis that CEP55 contributes to integrin-dependent oncogenic pathways and may represent a viable therapeutic target with minimal impact on normal adult tissues.

A key discovery in our study was the genetic interaction between Cep55 and the well-established tumor suppressor PTEN. While PTEN loss alone led to increased Akt signaling and promoted rapid development of tumorigenesis, simultaneous Cep55-deletion mitigated this phenotype. In the Cep55-PTEN double knockout (DKD), we observed delayed tumor onset, reduced tumor growth, and reduced downstream PI3K/Akt signalling compared to PTEN-KD alone. These findings suggest that Cep55 amplifies PI3K signaling and functions either downstream or parallel with PTEN, to contribute to tumor development and progression.

Moreover, spatial transcriptomics revealed that Cep55 deletion in the PTEN-null background induced immune-modulatory transcriptional programs, with upregulation of immune regulators (e.g., *Cd55*, *Dapl1*) and downregulation of immunosuppressive genes (e.g., *Serpine1*, *Cd207*). Enhanced expression of *Ptprc* (CD45) and B220 also indicated increased B-cell infiltration. Given the known immunosuppressive effects of PTEN loss including recruitment of myeloid-derived suppressor cells and T-cell exclusion, these findings suggest that CEP55 ablation may partially restore anti-tumor immunity. This opens potential for CEP55-targeted therapies to act synergistically with immunotherapy in PTEN-deficient cancers.

One limitation of our study was the rapid lymphoma onset and lethality in the PTEN-KD model, which restricted long-term mechanistic studies. As lymphoma represents the predominant and most lethal malignancy in this model, we focused on lymphoid organs for detailed analysis. Despite this constraint, the striking seven-fold survival extension in DKD mice enabled us to uncover later-onset, less aggressive tumors (e.g., papillomas, hemangioendotheliomas, prostate hyperplasia). While a broader analysis of tissue-specific tumor incidence would require conditional knockouts across organ systems, these findings reinforce the cancer-specific importance of CEP55, particularly in PTEN-deficient contexts. Sex differences also emerged: female DKD mice exhibited earlier mortality, often due to T-cell lymphomas, whereas 60% of males survived beyond 5 months, suggesting sex-specific influences on tumor biology that warrant further investigation.

Together, these findings position CEP55 as a context-dependent modulator of integrin trafficking, cell adhesion, and oncogenic signaling. Its dispensability in adult tissues and its synthetic vulnerability in PTEN-deficient tumors highlight it as a compelling therapeutic target. Future studies should explore CEP55 inhibition in preclinical models, assess combinatorial strategies with immunotherapy or PI3K inhibitors, and refine our understanding of the CEP55–integrin–PI3K axis in diverse cancer settings.

## Conclusion

This study identifies CEP55 as a promising, cancer-specific therapeutic target, particularly in PTEN-deficient tumors.. Through integrated proteomic, genetic, and spatial transcriptomic analyses, we show that CEP55 deletion impairs PI3K–AKT signaling, disrupts integrin trafficking, and enhances anti-tumor immunity. Importantly, normal adult tissues tolerate CEP55 loss, supporting its favorable therapeutic index. By prolonging survival and reducing tumor burden in a PTEN-null mouse model, CEP55 emerges as a multifunctional driver of malignancy and a potential vulnerability in aggressive cancers. Targeting CEP55 could provide a novel strategy to suppress tumor progression, particularly in PI3K/AKT-driven and immune-evasive cancers that are resistant to existing therapies.

## Abbreviation

CEP55: Centrosomal protein of 55 KD
cKO: Conditional KO (inducible)
DKD: Double knockdown
E1A/Ras: Adenovirus E1A + Rat sarcoma virus
ECM: Extracellular matrix
EV: Empty vector
GEMM: Genetically Enginered Mouse Models
Het: Heterozygous
Hom: Homozygous
IMM: immortalized
KD: Knock-down
KO: Knockout (constitutive)
MEFs: Mouse Embryonic Fibroblasts
OE: Overexpression
pMEFs: Primary MEFs
ROI: Region of interest
SV40: Simian virus 40 Large T Antigen
TMA: Tissue microarray
TME: Tumor microenvironment
WB: Western blot
Wt: Wild-type

## Acknowledgments

We thank the International Knockout Mouse Consortium and the Australian Phenomics Network Histopathology Facility at the University of Melbourne for providing valuable resources and support. We also acknowledge the QIMR Berghofer Medical Research Institute animal, histology, flow cytometry, and microscopy facilities, particularly Nigel Waterhouse and Tam Hong Nguyen, for their expert assistance.

Special thanks to Harsha Gowda and Rebekah Ziegman for performing the proteomics experiments.

## Authors’ contribution

Conceptualization: BR, PMA, KK; Investigation: BR, PT, PA, BB, BBW, SI, NJWvH, PMA; Bioinformatics: BR, YY, CV, SMT, BBW, ZX, HV, PHGD, PMA, QN; Supervisory team: ALB, JAL, BR, PMA, KK,; Histology and Pathology: SS, ZX, AS, JF; Data analysis: BR, BBW, AP, PMA; Writing the original draft: BR, ALB, PMA, KK; Review and Editing: BR, ST, PMA, KK; All authors read and approved the final manuscript.

## Data availability

Rashidieh, Ben; Ziegman, Rebekah; Khanna, Kum Kum (2025), “Genetic Ablation and Multi-Omics Profiling Reveal CEP55 as a Key Driver of Tumorigenesis in Diverse Cancer Models”, Mendeley Data, V1, doi: 10.17632/kkmmx9xnp4.1; 10.17632/ykffbkx4bs.1; 10.17632/49ggr7f2dc.1

## Extended Materials and Methods

### Animal Husbandry and Ethics Statement

Experimental animals utilized in this study were primarily C57BL/6J mice, except NOD/SCID mice employed in xenograft experiments. Mice were housed under controlled environmental conditions (25°C, 12-hour light/dark cycle) at the QIMR Berghofer Medical Research Animal Facility using OptiMICE® cages (Centennial, Colorado, USA). All experiments strictly complied with the Australian Code for the Care and Use of Animals for Scientific Purposes, approved by the QIMR Berghofer Animal Ethics Committee (approval number A0707-606M).

### Generation of Conditional Cep55 Knockout Mice

Cep55 floxed embryonic stem cells, obtained from the International Knockout Mouse Consortium, were used by the Australian Phenomics Network to produce heterozygous Cep55-targeted mice. These transgenic (Tg) mice possessed a “knockout-first” allele that facilitated constitutive Cep55 knockout or conditional allele generation upon further crossing. Conditional Cep55 knockout mice (Cep55 cKO) were generated by crossing heterozygous Cep55^Tg/+^ mice with FlpE mice to remove the neo cassette, resulting in Flp; Cep55^Fl/+^ progeny. Subsequent backcrossing with wild-type C57BL/6 mice removed the Flp transgene. Further intercrossing of Cep55^Fl/+^ mice produced Cep55Fl/Fl offspring. These mice were crossed with RosaCre^ERT2^ transgenic mice, generating RosaCre^ERT2^; Cep55^Fl/+^ mice, which were bred again with Cep55^Fl/Fl^ mice to obtain RosaCre^ERT2^; Cep55^Fl/Fl^ progeny. Tamoxifen administration activated Cre recombinase, resulting in Cep55 conditional knockout mice.

### Genotype Analysis

Genomic DNA, extracted from mouse ear samples using QuickExtract™ DNA Extraction Solution (Lucigen, USA), was used for genotyping. PCR employed a 3-primer strategy with a common forward primer (P1) and two distinct reverse primers (P2, P3). Wild-type mice produced a single band (381 bp), heterozygous mice two bands (381 bp, 454 bp), and homozygous knockout mice one band (454 bp). Primer sequences for Cre, FLPe, Pten, and Kras were previously published, while new sequences are listed: Cep55 (TGGGTCTTTAACTCATGGTC, AGGAGTGAAAAGTCCTCACA, GTACCGCGTCGAGAAGTT); Cre recombined (AACTGATGGCGAGCTCAGA); FLPe (GTGGATCGATCCTACCCCTTGCG, GGTCCAACTGCAGCCCAAGCTTCC); Pten (GCCCCGATGCAATAAATA, ACTCAAGGCAGGGATGAG); rcm2_CreF (TGTGGACAGAGGAGCCATAAC, CATCACTCGTTGCATCGACC); ROSA26 locus (AGCACTGGAAATGTTACCAAGGAAC, GGCTGGCTAAACTCTGGCCCTACA)

### Timed Mating

Successful timed mating was confirmed by observing copulation plugs the following morning, designated embryonic day 0.5 (E0.5).

### Organ and Embryo Isolation

Mice were anesthetized with Attane™ Isoflurane (Biomac Pty Ltd) and euthanized via cervical dislocation. Organs and embryos, dissected using a Nikon SMZ45 stereo microscope, were rinsed in ice-cold PBS. Samples were either snap-frozen for protein/mRNA extraction or fixed in Bouin’s solution or 4% paraformaldehyde (PFA) for histological analyses.

### Mouse cell derived transplant

Transplants were established in NOD/Scid mice by intraperitoneal injection of E1A/Ras-immortalized MEFs (1.0×10^6^ or 4.0×10^6^ cells suspended in Matrigel/PBS). Tumor growth was measured thrice weekly using calipers, and tumor volume calculated using the formula V=(W²×L)/2.

### Tamoxifen Induction

Cre recombination was induced *in vivo* using tamoxifen (1 mg) freshly prepared in sunflower oil/ethanol. MEFs underwent 4-hydroxy tamoxifen (4-OHT) treatment (1–3 µM) with optimal gene deletion observed at 1 µM after 72 hours.

### H&E and Immunohistochemistry staining

Tissues were fixed, embedded in paraffin, sectioned (4 µm), and processed for H&E and periodic acid–Schiff staining as previously described (Rashidieh et al., 2023). Antigen retrieval utilized sodium citrate buffer. Sections were permeabilized, blocked, incubated overnight with primary antibodies, and detected using Alexa Fluor–conjugated secondary antibodies. Images were acquired using Aperio® Scanscope® and analyzed via Image Scope and Imaris software.

### Cell Culture and MEF Establishment

Cell lines from ATCC were maintained per guidelines, authenticated by STR profiling, and tested annually for Mycoplasma. MEFs were isolated from e13 embryos. The MEFs were cultured in DMEM (20% FBS, antibiotics, antifungals) and immortalized via SV40T retroviral or E1A/Ras transduction.

### Cell Proliferation and Viability Assays

Proliferation was monitored using IncuCyte®. Viability was assessed by MTS assay, measuring absorbance at 490 nm post-treatment.

#### Clonogenic and Soft Agar Assays

MEFs seeded at low densities formed colonies after 14 days, quantified via crystal violet staining and imaging with GE InCell 2000 software.

### 3D Culture

MEFs cultured in Happy Cell® ASM medium formed spheroids, assessed by Hoechst staining and Cytell imaging after 14 days.

### Cell Invasion and Adhesion Assays

A custom invasion assay was established using a polyisocyanopeptide (PIC)-based hydrogel. PIC polymer hydrogel was synthesised with the collagen-mimicking peptide sequence Gly-Phe-Hyp-Gly-Glu-Arg (GFOGER) using previously described methods. PIC with a molecular weight Mv = 476 kg/mol containing 1% GFOGER ligand density (PIC-GFOGER) was dissolved in media to 2 mg/mL at 4 °C. To generate spheroids, cells were seeded in ultra-low attachment 96-well plates (Corning Costar CLS7007) at a density of 1,000 cells per well in complete DMEM medium. The cells formed spheroids of uniform size overnight and were maintained for 48 hours. After 48 hours, the spheroids were individually mixed with cooled PIC-GFOGER polymer solution on ice. The mixture was gently transferred to µ-Slide 15 Well 3D plates (Ibidi µ-Slide 15 Well 3D) and incubated at 37 °C to allow gelation. Thirty minutes after initial gelation, complete medium was added on top of the gel in each well. Spheroid outgrowth was monitored using a Leica SP8 confocal microscope with a 10× objective under brightfield settings. Spheroid diameters were measured using ImageJ. Adhesion assays involved collagen I-coated plates, crystal violet staining, and absorbance measurement at 570 nm.

### Live-Cell Imaging and Immunofluorescence

Live-cell imaging utilized EVOS Fl Auto and confocal microscopy. Immunofluorescence involved fixation, permeabilization, antibody staining, and imaging via DeltaVision microscopy.

### Western Blotting

Tissues and cells were lysed in RIPA or urea buffers, protein quantified via BCA assay, electrophoresed via SDS-PAGE, transferred to nitrocellulose membranes, and probed with antibodies listed: CEP55 CST (D1L4H) Rabbit mAb #81693 1:1000; Cep55 Santa Cruz biotechnology, sc-374051, 1:500; Vinculin Cell Signaling Technology #13901, 1:2000; β-Actin BD Pharmingen#612656, 1:2000; β-Catenin Cell Signaling Technology 9582, 1:1000; pERK1/2 (T202/Y204) Cell Signaling Technology #4370, 1:1000; ERK1/2Cell Signaling Technology #4695, 1:1000; pAKTS473 Cell Signaling Technology #4060, 1:1000; AKT Cell Signaling Technology #9271, 1:1000; MYC (Y69) Abcam #ab32072, 1:1000; Rabbit Secondary (peroxidase) Sigma Aldrich #A0545, 1:5000; Mouse Secondary (peroxidase) Sigma Aldrich #A9044, 1:5000.

### TCGA analyses

Messenger RNA levels, i.e., batch effect-normalized log2(normalized counts+1) (Illumina HiSeq RNASeqV2), were obtained from the pan-cancer dataset of The Cancer Genome Atlas (TCGA), including 33 cancer types (Weinstein et al., 2013) and processed as previously described (Shahrouzi et al., 2024).

### Immune cell infiltration levels in cancers

Estimation of the levels of 26 tumour-infiltrated immune cell types or states in TCGA samples occurred using CIBERSORTx, as previously described (Newman et al., 2019). These levels were compared to expression levels of CEP55 in the same samples using Spearman correlations. Spearman R and p values were determined in R (The R Project for Statistical Computing).

### Gene set enrichment analyses

For TCGA samples, single-sample Gene Set Enrichment Analysis (ssGSEA) was used as described (Barbie et al., 2009) to determine ssGSEA scores that reflect the activity of specific biological pathways or gene sets on a per-sample basis. The ssGSEA scores were compared to gene expression levels per sample. Heatmaps were created, with tiles showing the Spearman correlations (r and p values) between individual gene expression levels (CEP55 or PTEN) and the ssGSEA gene expression signatures per cancer type.

### Statistical Analysis

Analyses included Student’s t-test, ANOVA, Bonferroni post-tests, and log-rank tests using Prism v7.0 software. Significance was defined as: ns (P>0.05), * (P≤0.05), (P≤0.01), * (P≤0.001), and **** (P≤0.0001). Results were expressed as mean ± SD.

## Supplementary Tables

**ST1.**
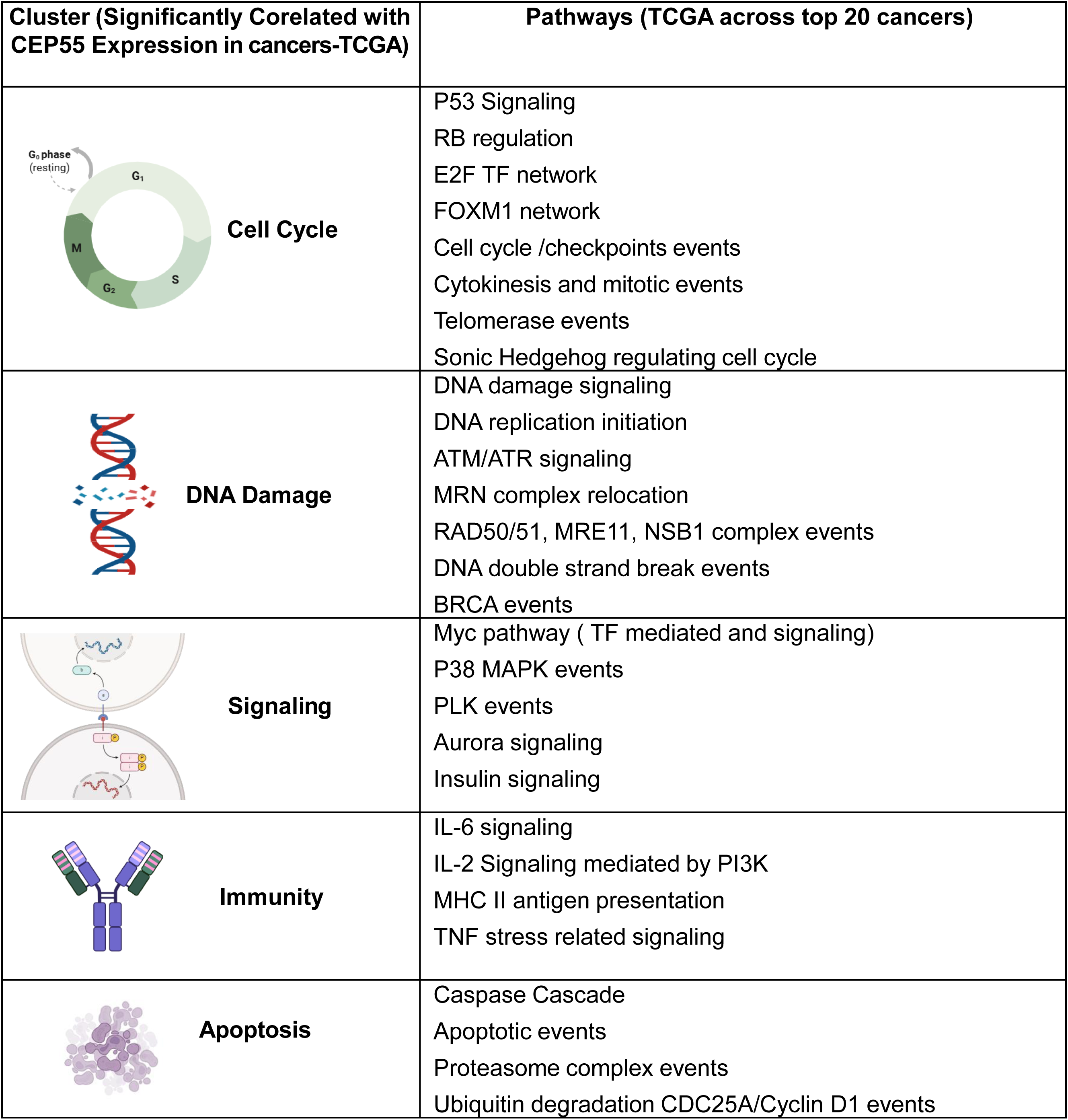
TCGA pathways correlated with CEP55 Expression.

| Cluster (Significantly Corelated with CEP55 Expression in cancers-TCGA) | Pathways (TCGA across top 20 cancers) |
| --- | --- |
| Cell Cycle | P53 Signaling<br>RB regulation<br>E2F TF network<br>FOXM1 network<br>Cell cycle /checkpoints events<br>Cytokinesis and mitotic events<br>Telomerase events<br>Sonic Hedgehog regulating cell cycle |
| DNA Damage | DNA damage signaling<br>DNA replication initiation<br>ATM/ATR signaling<br>MRN complex relocation<br>RAD50/51, MRE11, NSB1 complex events<br>DNA double strand break events<br>BRCA events |
| Signaling | Myc pathway ( TF mediated and signaling)<br>P38 MAPK events<br>PLK events<br>Aurora signaling<br>Insulin signaling |
| Immunity | IL-6 signaling<br>IL-2 Signaling mediated by PI3K<br>MHC II antigen presentation<br>TNF stress related signaling |
| Apoptosis | Caspase Cascade<br>Apoptotic events<br>Proteasome complex events<br>Ubiquitin degradation CDC25A/Cyclin D1 events |

**ST2.**
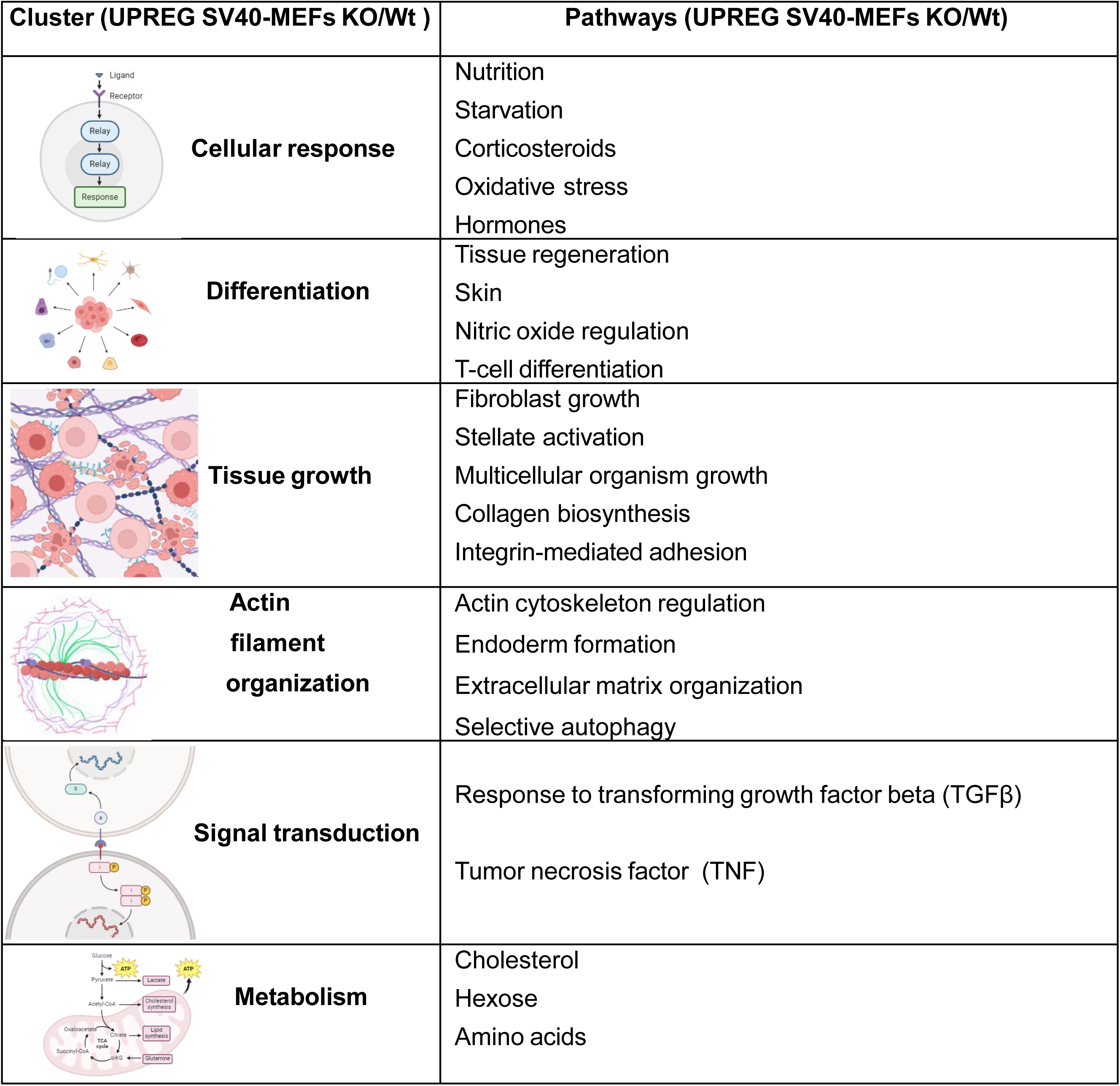
Differentially Upregulated pathways in Cep55 SV40-MEFS.

ST2. Differentially Upregulated pathways in Cep55 SV40-MEFs
| Cluster (UPREG SV40-MEFs KO/Wt ) | Pathways (UPREG SV40-MEFs KO/Wt) |
| --- | --- |
| Cellular response | Nutrition<br>Starvation<br>Corticosteroids<br>Oxidative stress<br>Hormones |
| Differentiation | Tissue regeneration<br>Skin<br>Nitric oxide regulation<br>T-cell differentiation |
| Tissue growth | Fibroblast growth<br>Stellate activation<br>Multicellular organism growth<br>Collagen biosynthesis<br>Integrin-mediated adhesion |
| Actin filament organization | Actin cytoskeleton regulation<br>Endoderm formation<br>Extracellular matrix organization<br>Selective autophagy |
| Signal transduction | Response to transforming growth factor beta (TGFβ)<br>Tumor necrosis factor (TNF) |
| Metabolism | Cholesterol<br>Hexose<br>Amino acids |

**ST3.**
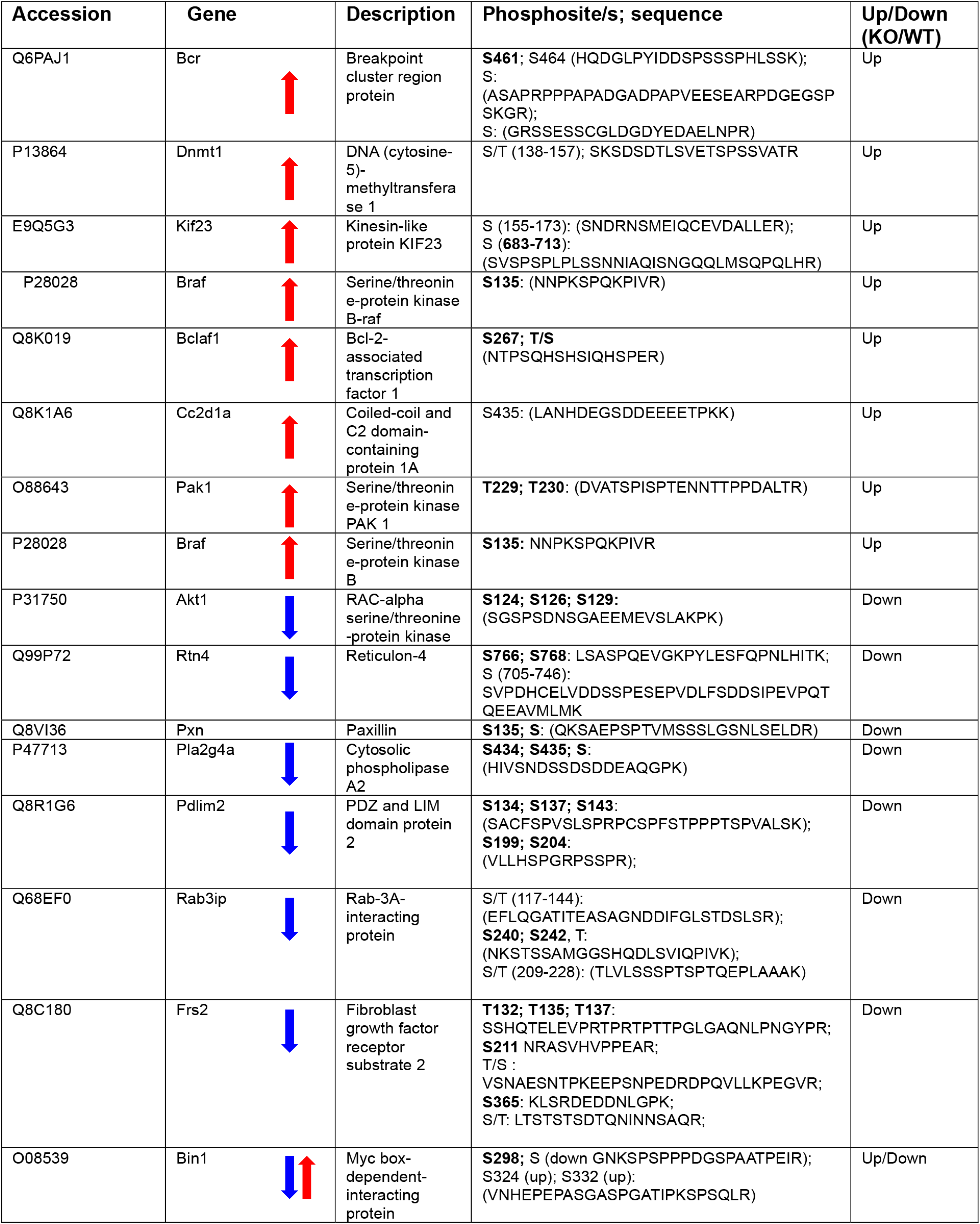
Differentially Up & downregulated phosphosites in E1A-RAS Cep55 KO MEFS.

**ST4.** Correlative Analysis of Indicated Gene Alterations & CEP55 Dysregulation.

| Query A | Query B | Log2 Odds Ratio | p-Value | q-Value | Tendency |
| --- | --- | --- | --- | --- | --- |
| ITGB1: AMP GAIN | CEP55: AMP GAIN | >3 | <0.001 | <0.001 | Co-occurrence |
| PTK2: AMP GAIN | CEP55: EXP >1.5 | 1.804 | 0.004 | 0.024 | Co-occurrence |
| ITGAV:H0MDEL HETLOSS | CEP55: EXP >1.5 | 2.057 | <0.001 | 0.007 | Co-occurrence |
| ITGA5:H0MDEL HETLOSS | CEP55: H0MDEL HETLOSS | 2.271 | <0.001 | <0.001 | Co-occurrence |
| ITGB1: HOMDEL HETLOSS | CEP55: H0MDEL HETLOSS | >3 | <0.001 | <0.001 | Co-occurrence |
| ITGAV= integrin αv<br>ITGA5= integrin α5<br>ITGB1= integrin β1<br>FN1: Fibronectin<br>PTK2: Focal adhesion kinase (FAK)<br>AMP/GAIN: Increased Copy number<br>EXP>1.5 mRNA expression is 1.5 SD above the mean<br>EXP<1.5 mRNA expression is 1.5 SD below the mean |  |  |  |  |  |

**ST5.** The observed pathological defects in Cep55-Pten DKD and Pten KD mice.

| Genotype | Cep55-Pten DKD | Pten KD |
| --- | --- | --- |
| Lymphoma | 11(16) | 6(8) |
| Extra-medullary hemopoiesis | 4(16) | 1(8) |
| Squamous papilloma | 4(16) | 0 |
| Hematoma | 5(16) | 1(8) |
| Hemangio-endothelioma (sarcoma) | 2(16) | 0 |

**ST6.**
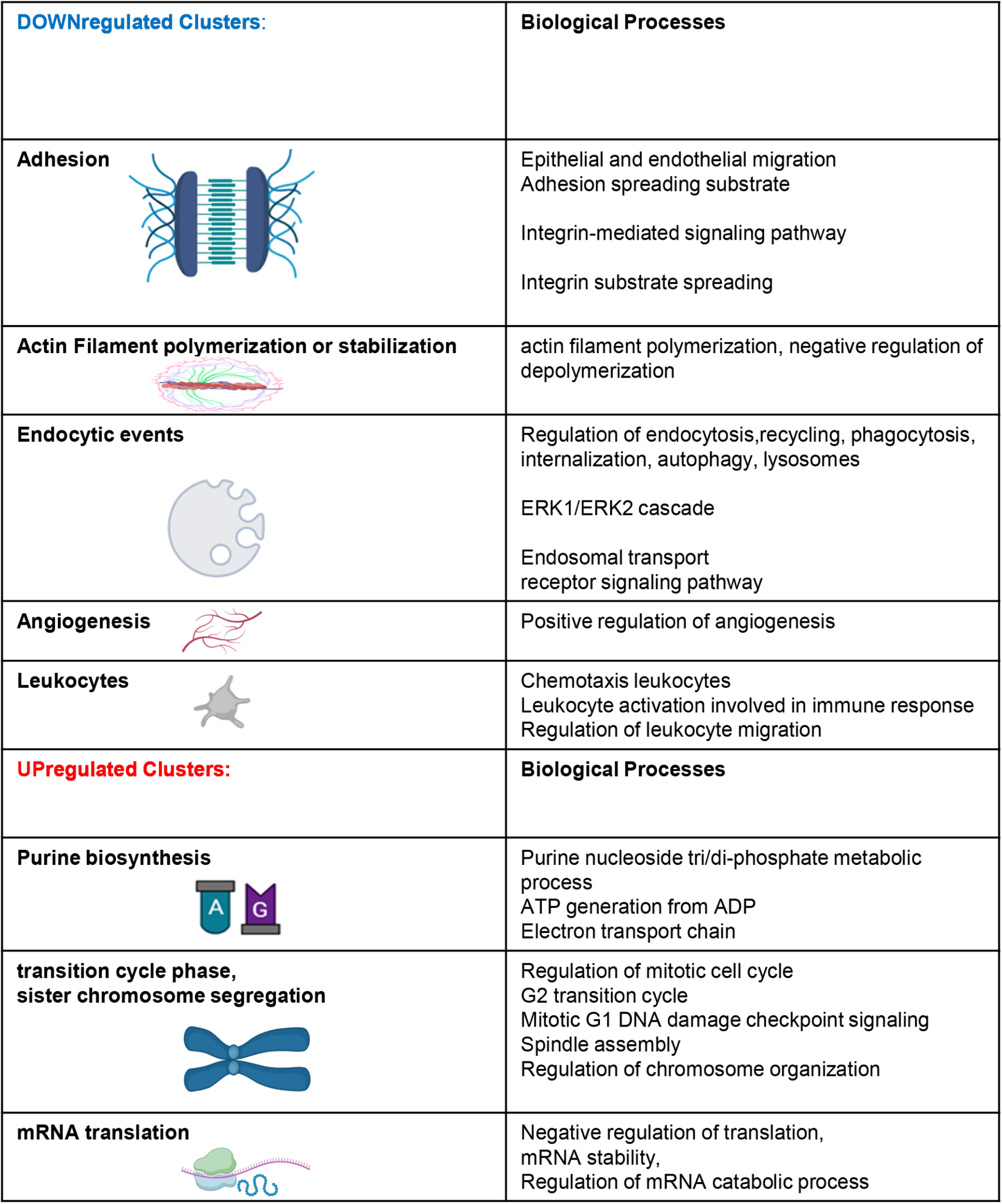
E1A/Ras KO MEFS & Cep55-Pten DKD.

|  |  |
| --- | --- |
| **DOWNregulated Clusters:** | **Biological Processes** |
| **Adhesion** | Epithelial and endothelial migration<br>Adhesion spreading substrate<br>Integrin-mediated signaling pathway<br>Integrin substrate spreading |
| **Actin Filament polymerization or stabilization** | actin filament polymerization, negative regulation of depolymerization |
| **Endocytic events** | Regulation of endocytosis,recycling, phagocytosis, internalization, autophagy, lysosomes<br>ERK1/ERK2 cascade<br>Endosomal transport receptor signaling pathway |
| **Angiogenesis** | Positive regulation of angiogenesis |
| **Leukocytes** | Chemotaxis leukocytes<br>Leukocyte activation involved in immune response<br>Regulation of leukocyte migration |
| **UPregulated Clusters:** | **Biological Processes** |
| **Purine biosynthesis** | Purine nucleoside tri/di-phosphate metabolic process<br>ATP generation from ADP<br>Electron transport chain |
| **transition cycle phase, sister chromosome segregation** | Regulation of mitotic cell cycle<br>G2 transition cycle<br>Mitotic G1 DNA damage checkpoint signaling<br>Spindle assembly<br>Regulation of chromosome organization |
| **mRNA translation** | Negative regulation of translation, mRNA stability, Regulation of mRNA catabolic process |

## Supplementary Figures

**Sup fig 1:**
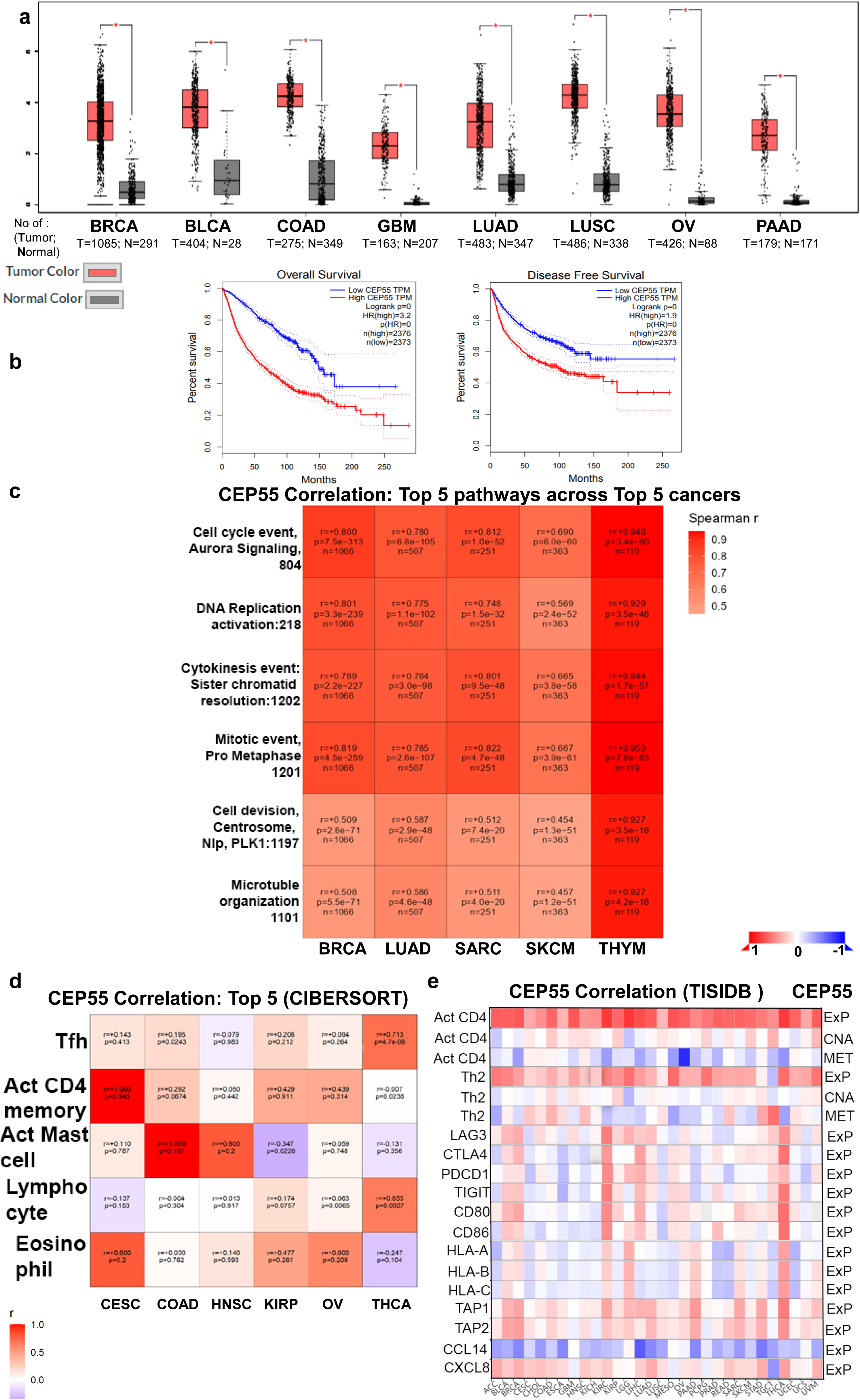

**Sup fig 2:**
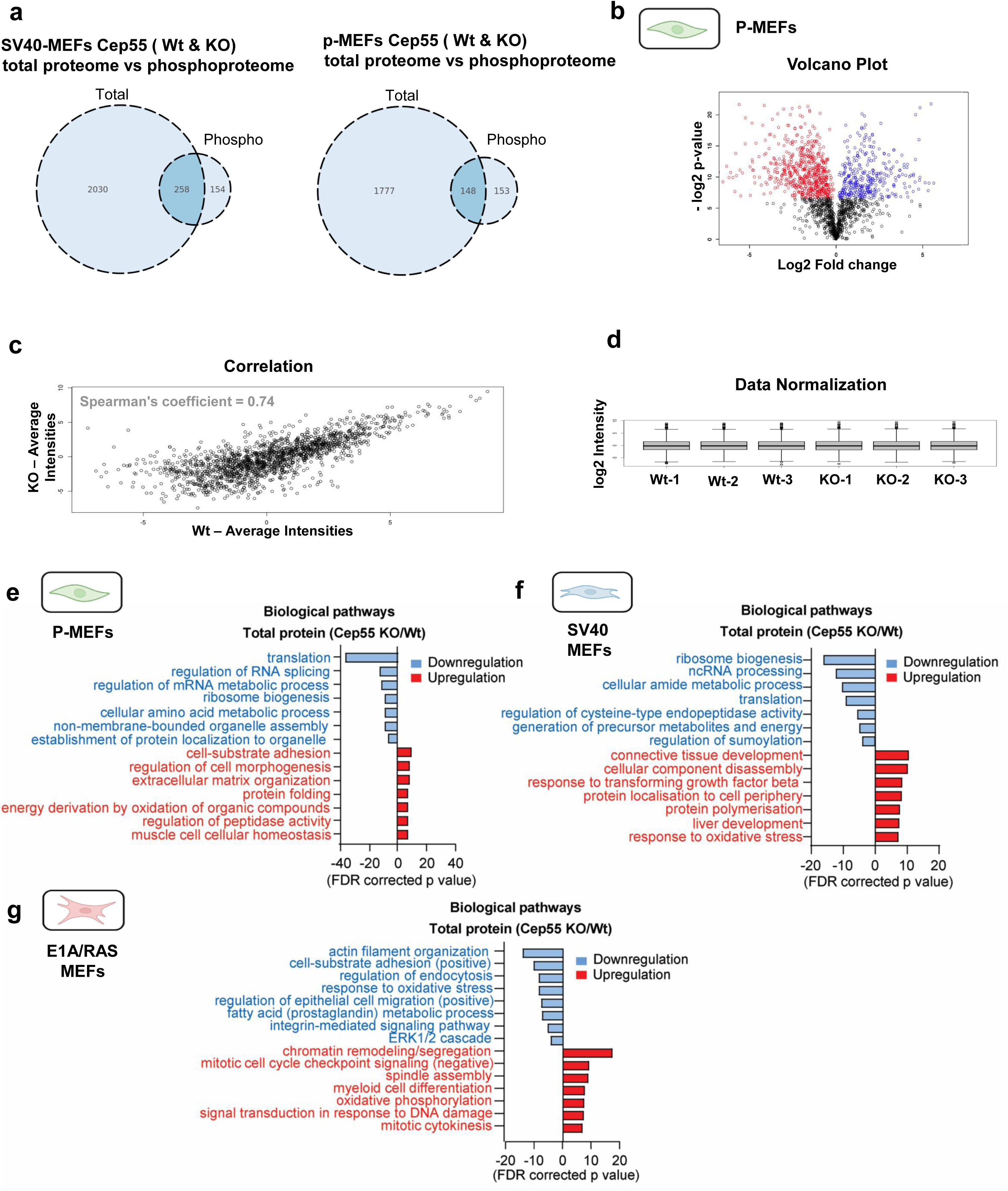

**Sup fig 3:**
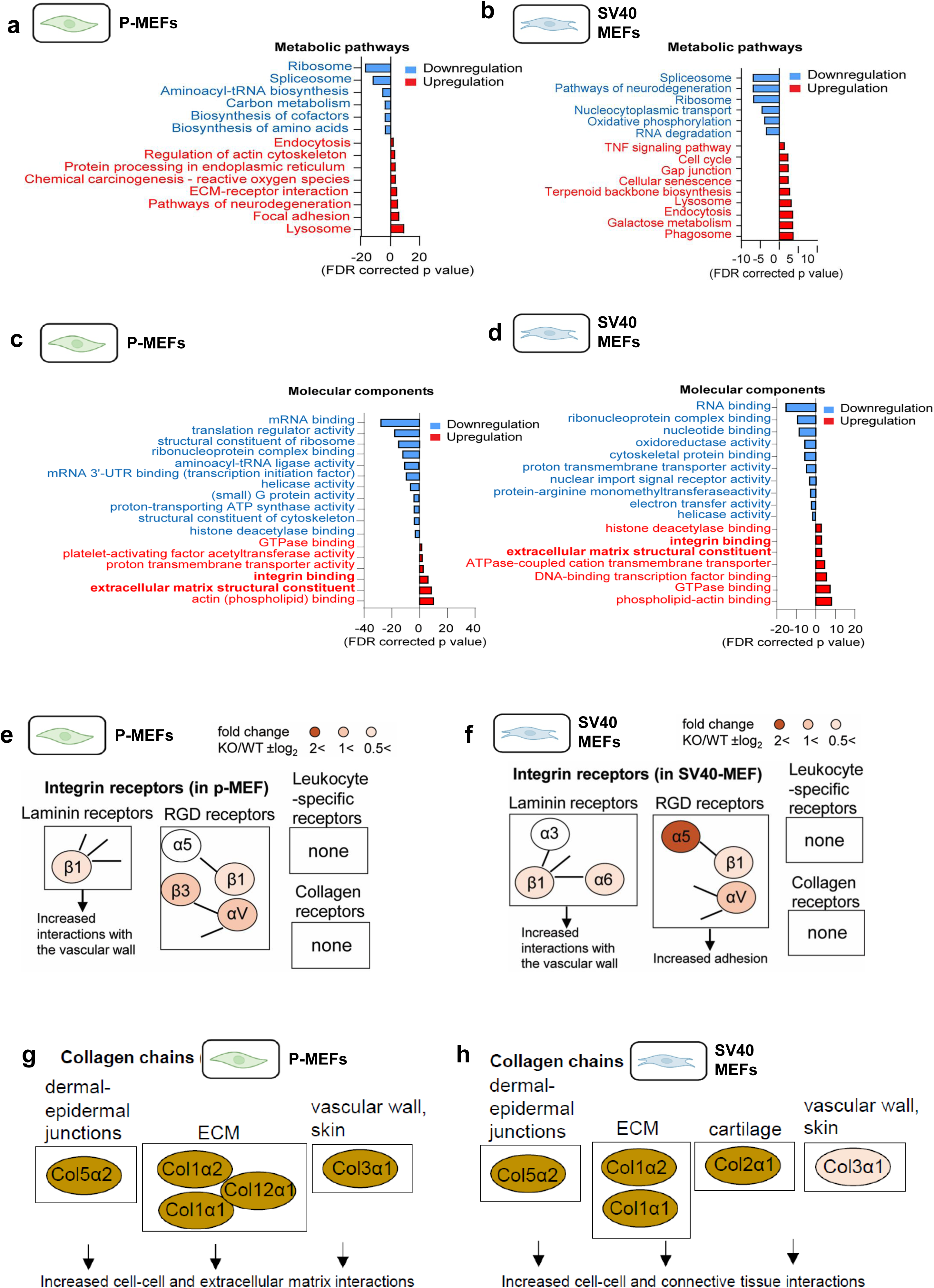

**Sup fig 4:**
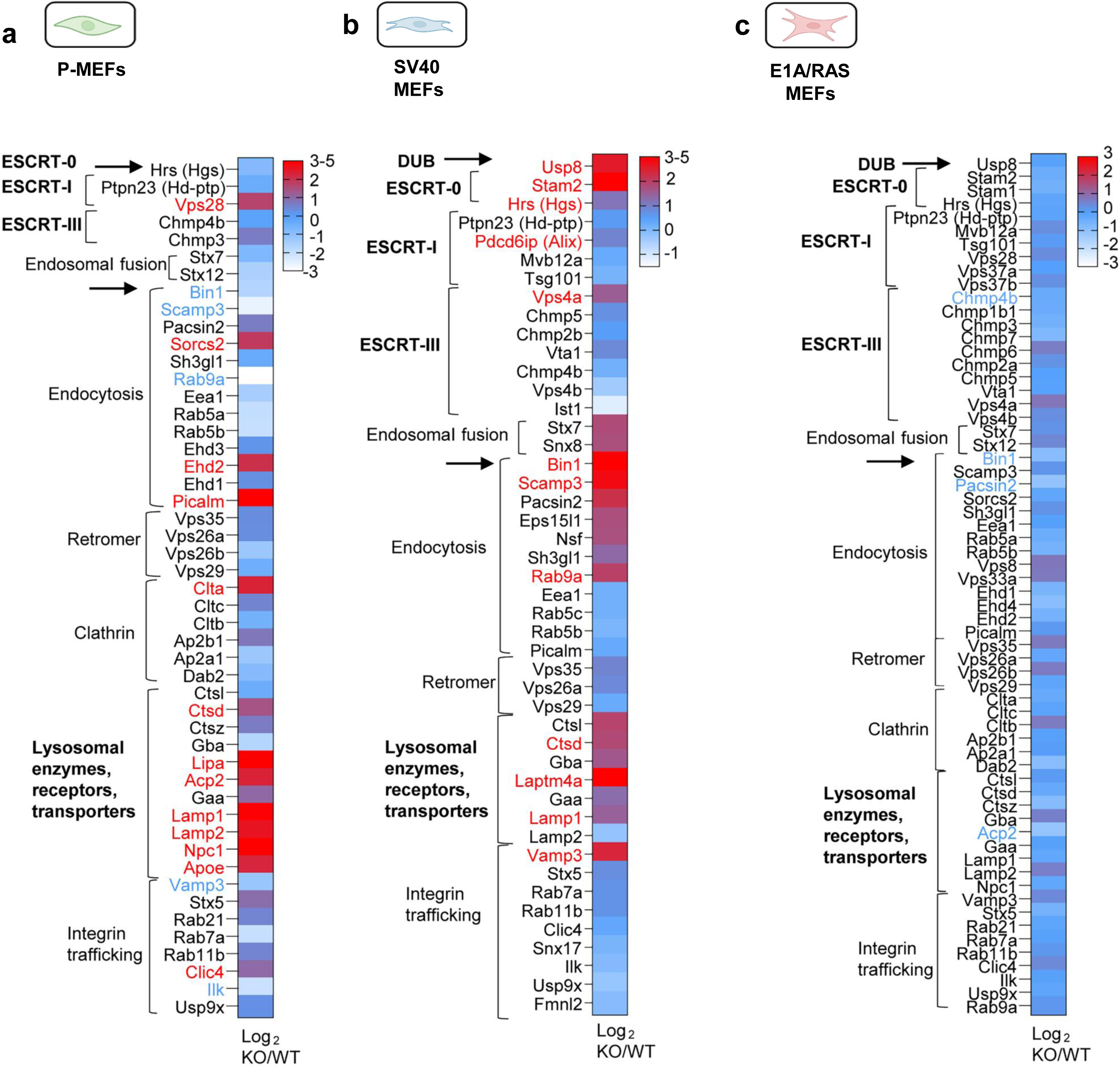

**Sup fig 5:**
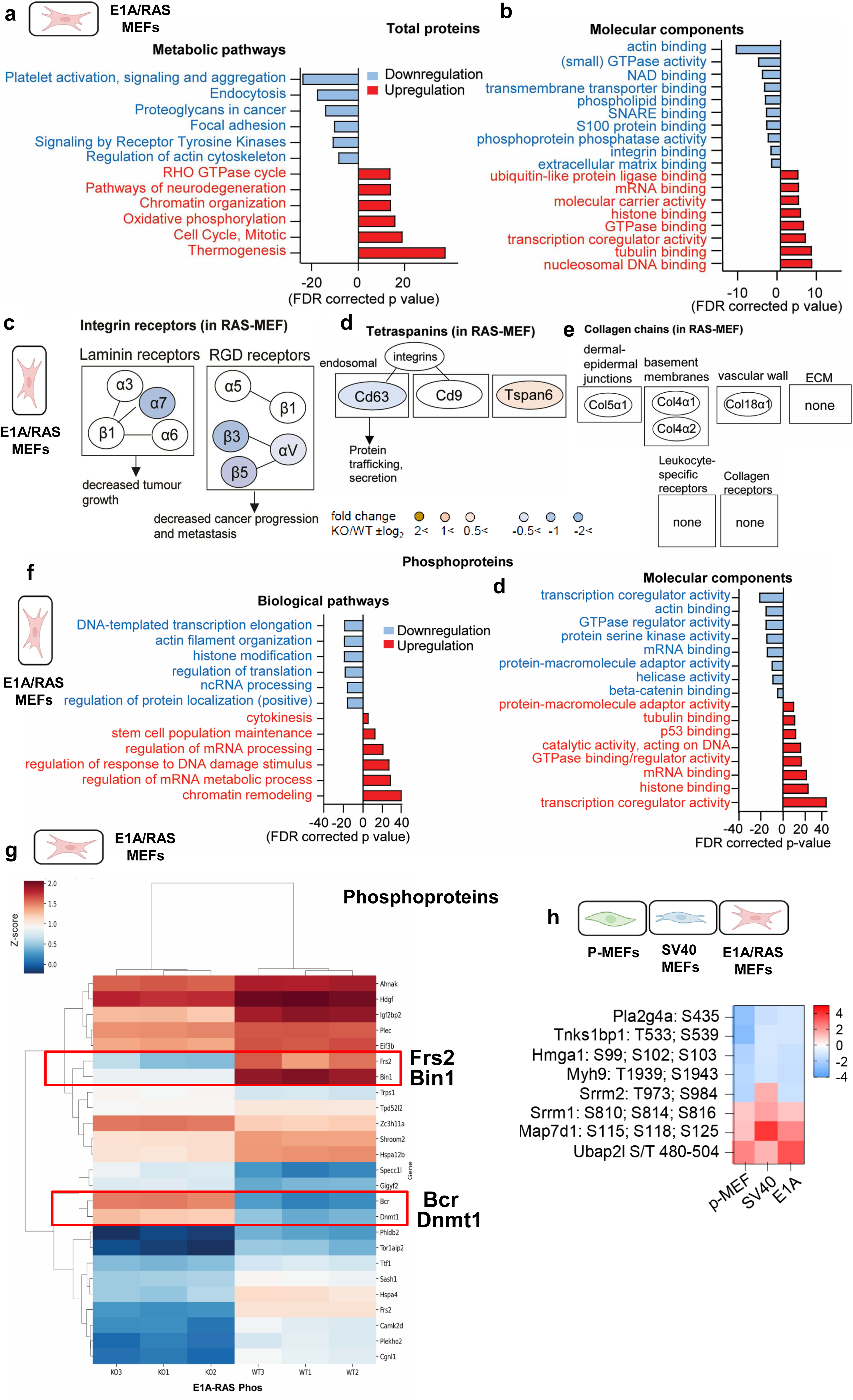

**Sup fig 6:**
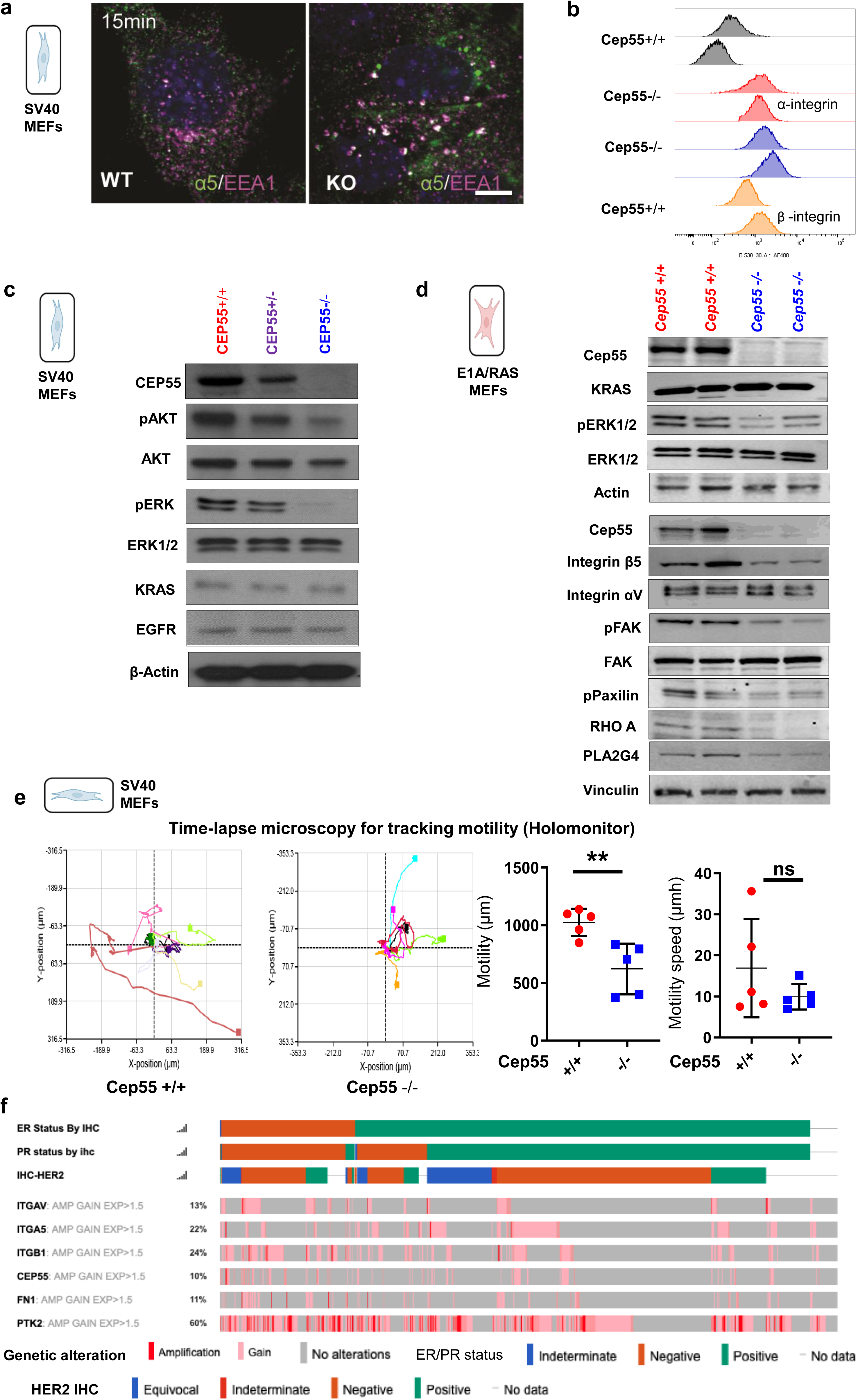

**Sup fig 7:**
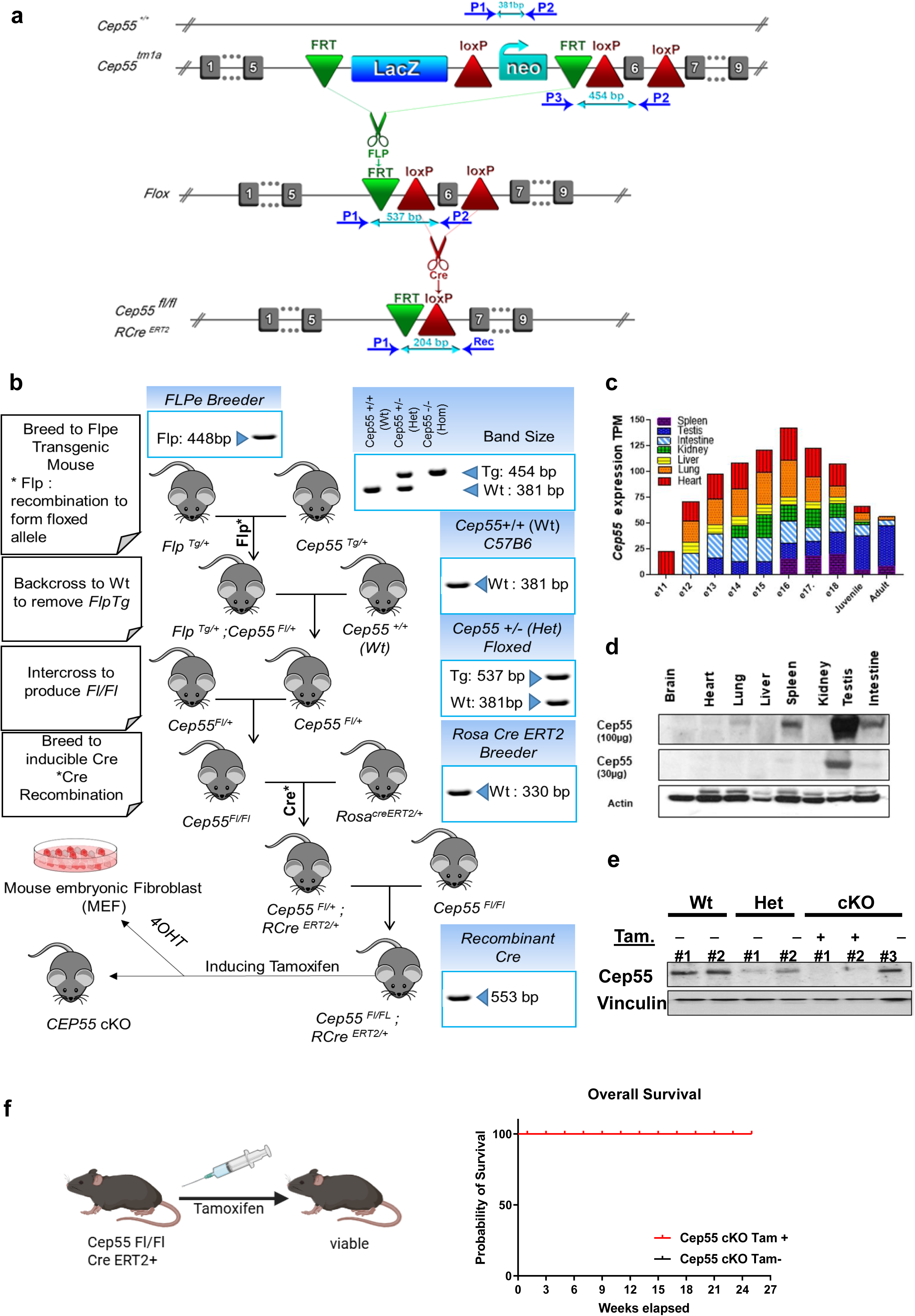

**Sup fig 8:**
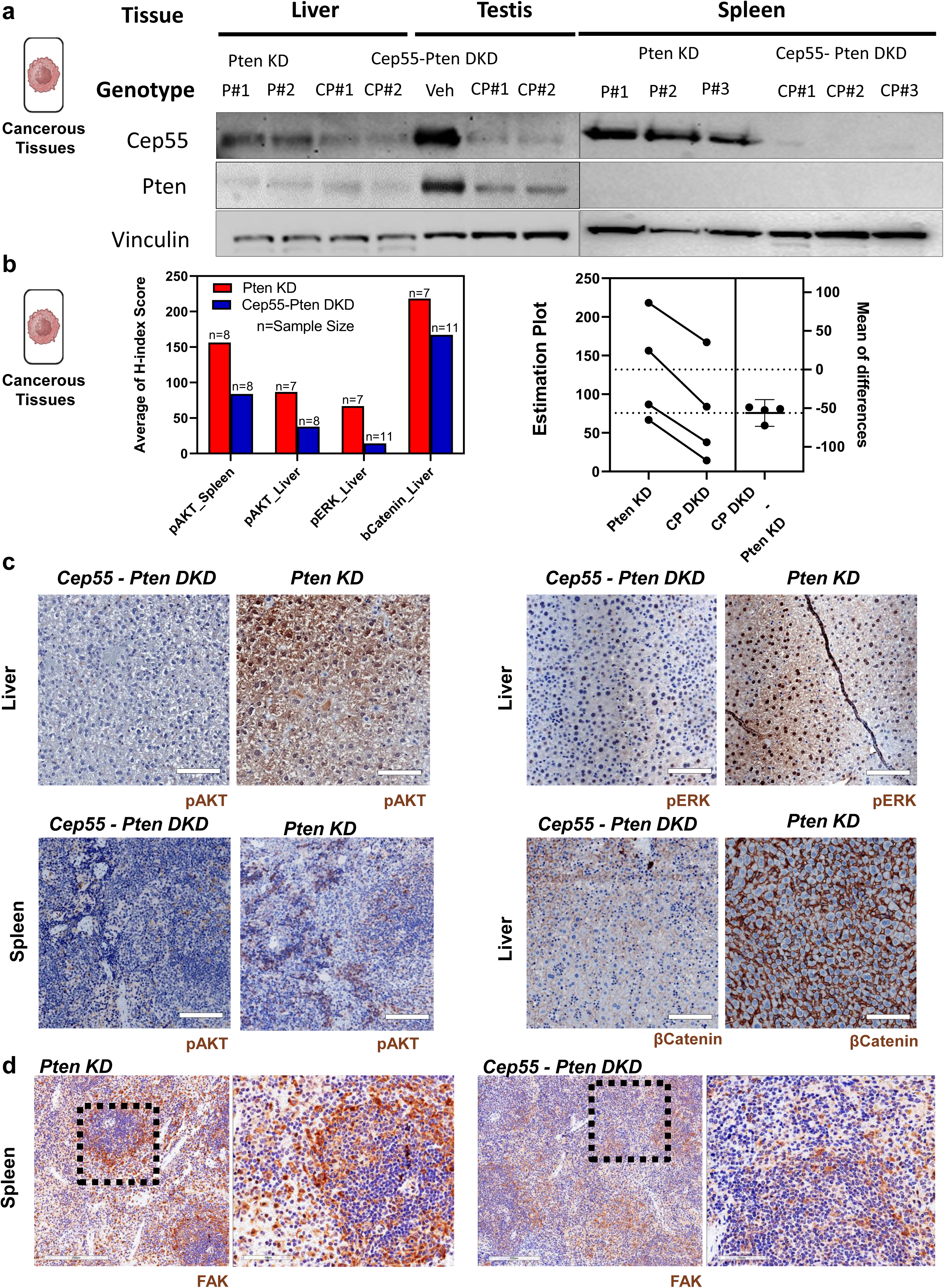

**Sup fig 9:**
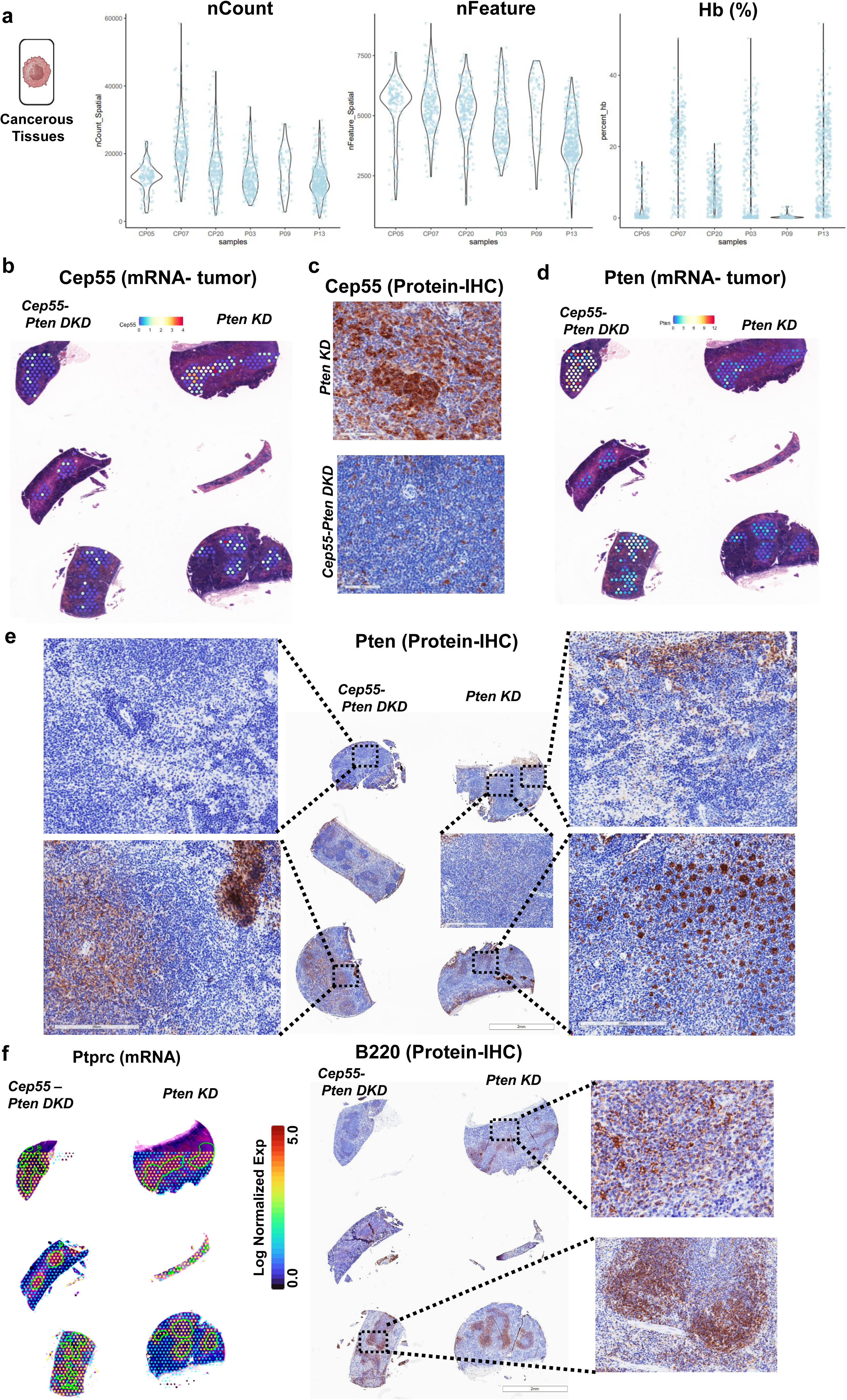

**Sup fig 10:**
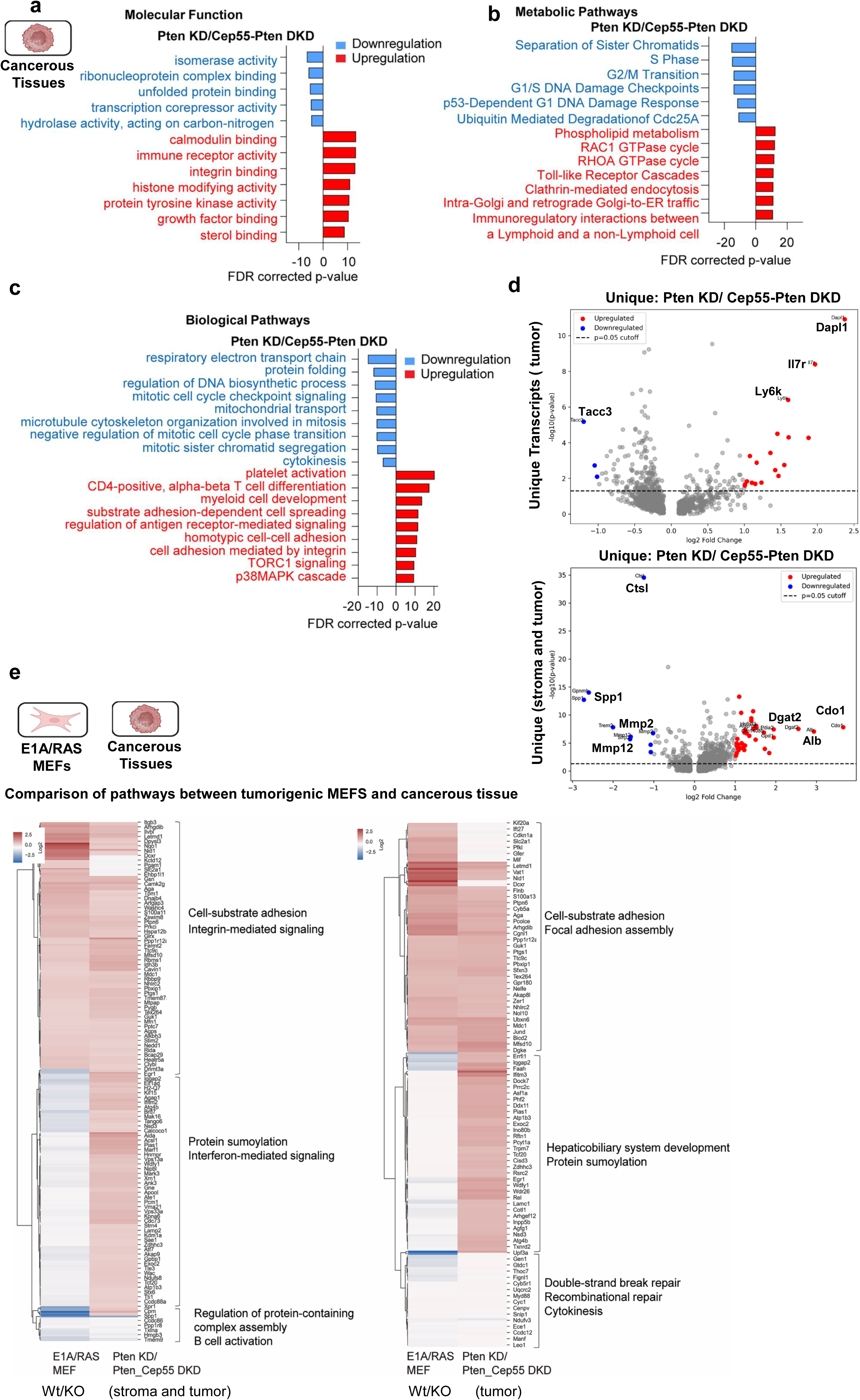

